# A Novel Role for FSTL-1 in Hematopoietic Stem Cell Homeostasis and Post-injury Regeneration

**DOI:** 10.64898/2026.10.07.757307

**Authors:** Craig A. Byersdorfer, Matthew Henkel, Norie Sugitani, Dequan Lou, Kong Chen, Brian T. Campfield

## Abstract

**Background:** The ability to repopulate all lineages of blood cells from a single hematopoietic stem cell (HSC) requires a constant balancing between maintenance of quiescence/self-renewal and the need for differentiation and adoption of specific lineage phenotypes. A classical interpretation suggests that during homeostasis, long- and then short-term HSCs differentiate into multipotent progenitors (MPPs), which then lack self-renewal capacity but can still differentiate into all downstream lineages. While much is known about the intrinsic and extrinsic regulators that promote HSC homeostasis, less is known about factors which influence HSC behavior following significant bone marrow (BM) injury, such as sublethal irradiation (SLI).

**Methods:** Conditional knock-out of follistatin-like 1 (FSTL-1) was achieved by breeding Fstl1 fl/fl mice to mice expressing CreERT2 under control of the ROSA locus, followed by systemic tamoxifen administration. ELISA and RT-PCR quantitated protein and RNA level loss of FSTL-1, respectively. HSC populations were assessed by flow cytometry under multiple conditions. Sublethal irradiation induced BM injury and blocking polyclonal FSTL-1 antibodies were used to recapitulate results seen with genetic ablation. Finally, single cell RNA sequencing defined both cell-specific expression of *Fstl1* within the BM compartment, but also identified transcripts and pathways upregulated in FSTL-1’s absence.

**Results:** These studies delineate a novel role for FSTL-1 in regulating HSC biology. In addition to selective expression of *Fstl1* within HSCs, genetic deletion of *Fstl1* increases Lin−Sca-1+c-Kit+ (LSK) cell number and alters HSC homeostasis. These findings are further amplified in the BM in the context of recovery from sublethal irradiation. Importantly, the post-irradiation increase in BM cellularity seen with *Fstl1* deletion is replicated by treatment with anti-FSTL-1 polyclonal antibodies. Finally, single cell RNA sequencing reveals distinct *Fstl1* expression in specific LSK subpopulations, including HSCs and proliferating cells, and identifies a novel relationship between decreased FSTL-1 levels and enrichment of canonical metabolic pathways including oxidative phosphorylation, fatty acid metabolism, and glycolysis.

**Conclusions:** Taken together, these results highlight a novel role for FSTL-1 in promoting HSC homeostasis. They also unlock an innovative way to promote HSPC recovery following BM injury, knowledge predicted to be of significant therapeutic value.

## Background

Hematopoietic stem cells (HSCs) are responsible for generating all mature hematopoietic lineages, including red blood cells, white blood cells, and platelets. Long-term (LT)-HSCs are at the top of this differentiation hierarchy and must maintain a constant balance between maintenance of quiescence and self-renewal capacity while still producing sufficient downstream progenitors in response to the hematopoietic needs of the organism. Slightly more differentiated multipotent progenitors (MPPs), in contrast, offer limited self-renewal capacity, yet remain capable of differentiating into all lineages of hematopoietic cells, leading to the paradigm that during homeostasis, LT- and then short-term (ST)-HSCs repopulate MPP pools, which in turn maintain blood cell populations over the course of a lifetime [1, 2]. Along the way, disruptions such as inflammatory signaling, aging-associated stress, and altered niche composition can impair MPP generation and thus skew differentiation, yielding dysfunctional hematopoiesis [3]. Further, injury to the hematopoietic niche (e.g. chemotherapy, irradiation, or sepsis) favors a more prominent role for MPP cells, with only a fraction of potential HSC clones differentiating into blood cell production [4, 5]. Finally, despite decades of work on understanding HSC biology, a complete delineation of all the factors which govern decisions during the process of stress-induced hematopoiesis remains elusive.

The maintenance of HSC quiescence is an active process, sustained by actions of both cell intrinsic regulators and cell extrinsic (i.e., niche-derived) signals. These regulators and signals work together during homeostasis to limit HSC proliferation and preserve LT-HSC self-renewal. Key intrinsic factors include p27 and p57 [6, 7], transcriptional regulators from the FOXO family of proteins [8], p21 [9], and the metabolic regulator mammalian target of rapamycin [10]. Together, these factors enforce cell-cycle arrest [11, 12] and maintain minimal levels of metabolic activity, predominated by a low degree of mitochondrial respiration and a preferred reliance on glycolysis [13]. Extrinsically, the BM niche provides signals such as TGF-β and CXCL12 to reinforce the HSC quiescence by suppressing cell cycling and promoting stem cell retention [14–16]. Interestingly, although many factors responsible for promoting LT-HSC quiescence have been described, less is known about factors whose absence increases stem cell differentiation.

Follistatin-like 1 (FSTL-1) is a 35 kilodalton secreted glycoprotein, with two O- and four N-glycosylation sites. It exerts a myriad of functions in mammalian biology, including specific roles in embryogenesis, inflammation, and tissue repair [17–24]. Specifically, FSTL-1 promotes cardiac regeneration via regulation of cell-cycling [25], enhances regenerative potential in mesenchymal stem cells (MSCs) [26], and promotes stemness in breast cancer cells [27]. Importantly, our group recently described a novel role for FSTL-1 in promoting expression of nuclear receptor subfamily 4 group A member 1 (Nr4a1) and Krüppel-like factor 4 (Klf4) in lung tissue [28]. Notably, Nr4a1 and Klf4 are known regulators of HSC quiescence and self-renewal [29–31]. From the standpoint of regulation, post-translational modifications play an important role in FSTL-1 function [32], with subsequent glycosylation being an important determinant of its overall biological activity [33].

Here we describe a novel role for FSTL-1 in governing HSC biology. Genetic loss of FSTL-1 increases the percentage and total number of Lineage negative, Sca-1+c-Kit+ (LSK) cells recovered during BM homeostasis, a finding that is further amplified during recovery from sublethal irradiation (SLI). Furthermore, these increases in cellularity following irradiation-induced injury could be replicated by inhibiting FSTL-1 function via antibody blockade, both *in vitro* and *in vivo*. Finally, single cell RNA sequencing defined expression of *Fstl1* in specific LSK subpopulations, while gene set enrichment analysis (GSEA) identified pronounced changes in metabolism- and proliferation-related gene pathways in Fstl1-deficient cells. Together, these results both support a novel role for FSTL-1 in governing HSC biology during BM homeostasis, but also identify an approach to enhance the recovery HSPCs following significant BM injury, findings that could be leveraged to powerful therapeutic advantage.

## Methods

### Mice

Mice were generated for conditional elimination of *Fstl1* by crossing the B6.129-Gt(ROSA)26Sor^tm1(cre/ERT2)Tyj^/J (Jackson Laboratories #:008463) strain [34] with mice homozygous for loxP sequences inserted into exon 1 and intron 2 of the *Fstl1* gene (FSTL-1^fl/fl^) (full generation detailed in [17]), with *Fstl1*-floxed mice a generous gift from Raphael Hirsch. All mice were housed in a University of Pittsburgh pathogen-free facility in accordance with the Institutional Animal Care and Use Committee (IACUC) established guidelines. Experiments were performed under the University of Pittsburgh’s IACUC approved protocols. Female and male mice (between 8- and 20-weeks-old) were used interchangeably for experiments.

### Tamoxifen administration and sublethal irradiation (SLI)

To ensure Cre translocation to the nucleus, mice were treated with 4mg tamoxifen in corn oil daily for two days via intraperitoneal (i.p.) injection, rested a day, then given two additional days of i.p. injections. Cre^neg^ (FSTL-1^fl/fl^) animals also received tamoxifen as a negative control. For homeostasis experiments, BM analysis was performed at 48 hours, 7 days, and 14 days from the first tamoxifen treatment. For SLI experiments, tamoxifen administration was followed on day 14 by 450 cGy of total body irradiation with a Precision MultiRad350. BM cells were collected 14 days post-irradiation for further analysis.

### Anti-FSTL-1 antibody

Purified MBP-tagged recombinant FSTL-1 was used as an immunogen to elicit rabbit polyclonal serum, which was Protein A column purified to isolate polyclonal, rabbit anti-mouse FSTL-1 antibodies (Biosynthesis, Lewisville, TX). Anti-FSTL1 was verified to be endotoxin-free prior to experimental treatment. For anti-FSTL-1 neutralizing antibody experiments *in vitro,* 2 million whole BM cells were incubated in 96-well round-bottom plates with anti-FSTL1 or control polyclonal IgG (R&D Systems #AB105C) at a concentration of 290ug/ml in complete DMEM and cultured for 24 hours prior to analysis. For *in vivo* studies, mice were treated with 50ug of anti-FSTL-1 or polyclonal IgG, in 200ul sterile PBS, i.p. on days -1, 0, +2 and then every other day, for three days a week, until study completion on Day 14 post-SLI (as described above).

### Bone Marrow Cell Isolation

Two femurs per mouse were cut at both ends, placed into a 0.6mL tube with a hole in the bottom, which was then placed into a 1.5mL tube and spun at 1200xg for 5 minutes at room temperature. Cell pellets were immediately resuspended in complete DMEM media containing 4.5g/L D-glucose (Gibco), 10% heat-inactivated fetal bovine serum (FBS, Gibco), 2mM L-glutamine (Gibco), 100U/mL penicillin, 100μg/mL streptomycin (Gibco), 1X non-essential amino acids (Gibco), 1mM sodium pyruvate (Gibco), and 10mM HEPES (Gibco). Single cell suspensions were then passed through a 70micron filter and washed twice with complete DMEM before a final resuspension in 1mL complete DMEM. Cells were enumerated on a Countess 3FL Automated Cell Counter using Trypan blue (Invitrogen).

### Flow Cytometry

Bone marrow single cell suspensions were checked for cell viability by incubating with LIVE/DEAD™ Fixable Aqua Dead Cell Stain Kit (Invitrogen) according to manufacturer’s instructions for 30 minutes on ice. When CD16/32 (clone 93) expression was not being analyzed within a given panel, cells were incubated with Fc block (eBioscience/Invitrogen) at a concentration of 1:100 in 2% heat-inactivated FBS-PBS solution for 15 minutes on ice. Surface antibodies were then incubated at a 1:100 concentration in 2% heat-inactivated FBS-PBS solution for 30 minutes on ice. Anti-mouse surface antibodies used in the present study include: Hematopoietic Lineage Antibody Cocktail (clones 17A2, RA3-6B2, M1/70, TER-119, RB6-8C5, FITC; eBioscience), anti-Sca-1 (clone D7, BV421; BD Biosciences), anti-CD117/c-Kit (clone 2B8, BV785; BioLegend), anti-CD48 (clone HM48-1, APC; BioLegend), anti-CD150 (clone TC15-12F12.2, PE; BioLegend), anti-CD16/32 (clone 93, PerCP-Cy5.5; BioLegend), anti-CD127/IL-7R (clone SB/199, PE-Cy7; BD), anti-CD4 (clone RM4-5, BV786; BD), anti-CD8a (clone 53-6.7, BV605; BioLegend), anti-CD19 (clone 1D3/CD19, PerCP-Cy5.5; BioLegend), anti-CD115 (clone AFS98, PE; BioLegend), anti-Ly-6G/Ly-6C/Gr-1 (clone RB6-8C5, FITC; BioLegend), anti-TCRβ (clone H57-597; APC-eFluor780; eBioscience/Invitrogen), anti-TER-119 (clone TER-119, BV421; eBioscience/Invitrogen). Data was acquired on a BD LSRFortessa (BD Biosciences) using FACS Diva Software v8.0.1 and analyzed using FlowJo software v10.9.0 (BD Biosciences).

### Mouse FSTL-1 ELISA

Serum samples were collected from mice via a terminal, retroorbital bleed. Blood was collected in a 1.5mL microcentrifuge tube, allowed to coagulate for 20 minutes at room temperature, and spun at 5,000xG for 15 minutes. The serum layer was removed to a new cryovial and banked at -80C storage until the time of ELISA. ELISAs were performed in flat-bottomed 96-well maxisorp plate (Nunc). MAB1738 (monoclonal rat anti-mFSTL1 IgG2a, R&D Systems) was used as the coating antibody at a final concentration of 5ug/mL per well in PBS and incubated at 4°C overnight. Following each incubation, unless specified otherwise, plates were washed 5 times with 1X ELISA/ELISPOT Assay Diluent (eBioscience/Invitrogen) to maximum well volume. The plate was blocked with 1X ELISA/ELISPOT Assay Diluent for 1 hour at room temperature. Recombinant mouse FSTL-1 (R&D Systems) was used to generate an FSTL-1 specific standard curve with the top dilution of 12,500pg/mL. Serum samples were diluted five-fold in 1X ELISA/ELISPOT Assay Diluent. Standards and diluted serum samples were incubated for 1 hour at room temperature. Biotinylated AF1738 (polyclonal goat anti-mFSTL1 IgG, R&D Systems) was used as the detection antibody at a final concentration of 0.5ug/mL per well in 1X ELISA/ELISPOT Assay Diluent and incubated for 1 hour at room temperature. Avidin-HRP (eBioscience) is added to the wells at a concentration of 1:250 in 1X ELISA/ELISPOT Assay Diluent and incubate for 20 minutes at room temperature. BD OptEIA TMB Substrate Reagent set (BD) was used per the manufacturer’s recommendations, and without a wash step in between, 1M sulfuric acid was used to stop the reaction and the plate was read at a wavelength of 450nm on a BIOTEK Synergy H1 Hybrid Reader using Gen5 v3.15 software (Agilent).

### Quantitative Real-Time PCR

Bone marrow single cell suspensions were pelleted at 500xG for 5 minutes at 4°C and pellets resuspended in RLTPlus buffer supplemented with beta-mercaptoethanol for RNA extraction using Qiagen’s RNeasy Plus Mini Kit (Qiagen) according to the manufacturer’s protocol. RNA was reverse transcribed into cDNA using iScript Reverse Transcription Supermix (Bio-Rad) according to the manufacture’s protocol. TaqMan Universal PCR Master Mix (Applied Biosystems) was used according to manufacturer specifications with primer assays for Hprt1 (Mm03024075) and Fstl1 (Mm00433371) and run on a Bio-Rad CFX96 Real-Time System C1000 Touch Thermal Cycler (Bio-Rad) using Bio-Rad CFX Maestro software v2.3 (Bio-Rad).

### Single cell RNA Sequencing

FSTL1-cKO (Cre^pos^) and Cre^neg^ littermates had BM harvested 48 hours following Tamoxifen initiation, followed by preparation of a single cell suspension and antibody staining for immediate fluorescence activated cell sorting of BM LSK cells. 10,000 LSK cells per animal underwent subsequent processing for single-cell RNA sequencing. Single-cell suspensions were processed using the 10x Genomics Chromium 5ʹ Gene Expression platform. Reads were aligned to mm10/GRCm38 and quantified with Cell Ranger (v7.0) and matrices imported into R (Read10X). Samples were multiplexed by cell hashing; HTO counts were CLR-normalized and demultiplexed (HTODemux, positive.quantile = 0.99), retaining singlets. In Seurat (v5.2.0), cells with 200–6,000 genes, <5% mitochondrial reads, and with genes detected in ≥3 cells were retained. Data were log-normalized, scaled, and analyzed by PCA; the first 15 PCs were used for SNN graph construction, Louvain clustering (resolution = 0.2), and UMAP visualization. Cell types were annotated by canonical markers. Marker genes were identified (FindAllMarkers, Wilcoxon), and subsets were re-clustered. Differential expression was performed using MAST. GSEA (fgsea, MSigDB hallmark, category H) ranked genes by average log₂ fold change.

### Data Analysis

Graphing and statistical analysis was performed using GraphPad Prism for Windows (version 10.0., San Diego, CA; www.graphpad.com). Unless directly state otherwise, all data are displayed as dot plots, where each individual dot represents a separate, biological replicate. Lines are drawn at the mean value. Unpaired two-tailed Student T test was used to determine statistical significance between groups. In all cases, *p<0.05, **p<0.01, ***p<0.001, and ****p<0.0001.

## Results

### FSTL-1 levels are dynamically regulated during bone marrow transplantation

Previous literature has suggested a role for FSTL-1 in the biology of HSPCs [35]. To explore this idea further, we first queried the BloodSpot database for FSTL-1 expression in cells of the hematopoietic system [36]. Searching under ‘mouse normal hematopoietic system’ in the BloodSpot dataset revealed preferential transcription of *Fstl1* in both LT- and ST-HSCs, as well as in lymphoid-primed multipotential progenitors (Figure 1A). We next evaluated the dynamics of FSTL-1 protein expression over the course of a standard bone marrow transplant. CD45.2+ C57Bl/6 animals were lethally irradiated and transplanted one day later with 5e6 whole BM cells from a congenic CD45.1+ animal. Serum was collected pre-transplant on day -7, on day 0 (the day of cell injection), and then on days +11 and +28 post-transplant. Serum levels of FSTL-1 increased by 45% at the time of transplant on day 0, with a subsequent return back to pre-transplant levels by day +11 (Figure 1B).

**Figure 1:**
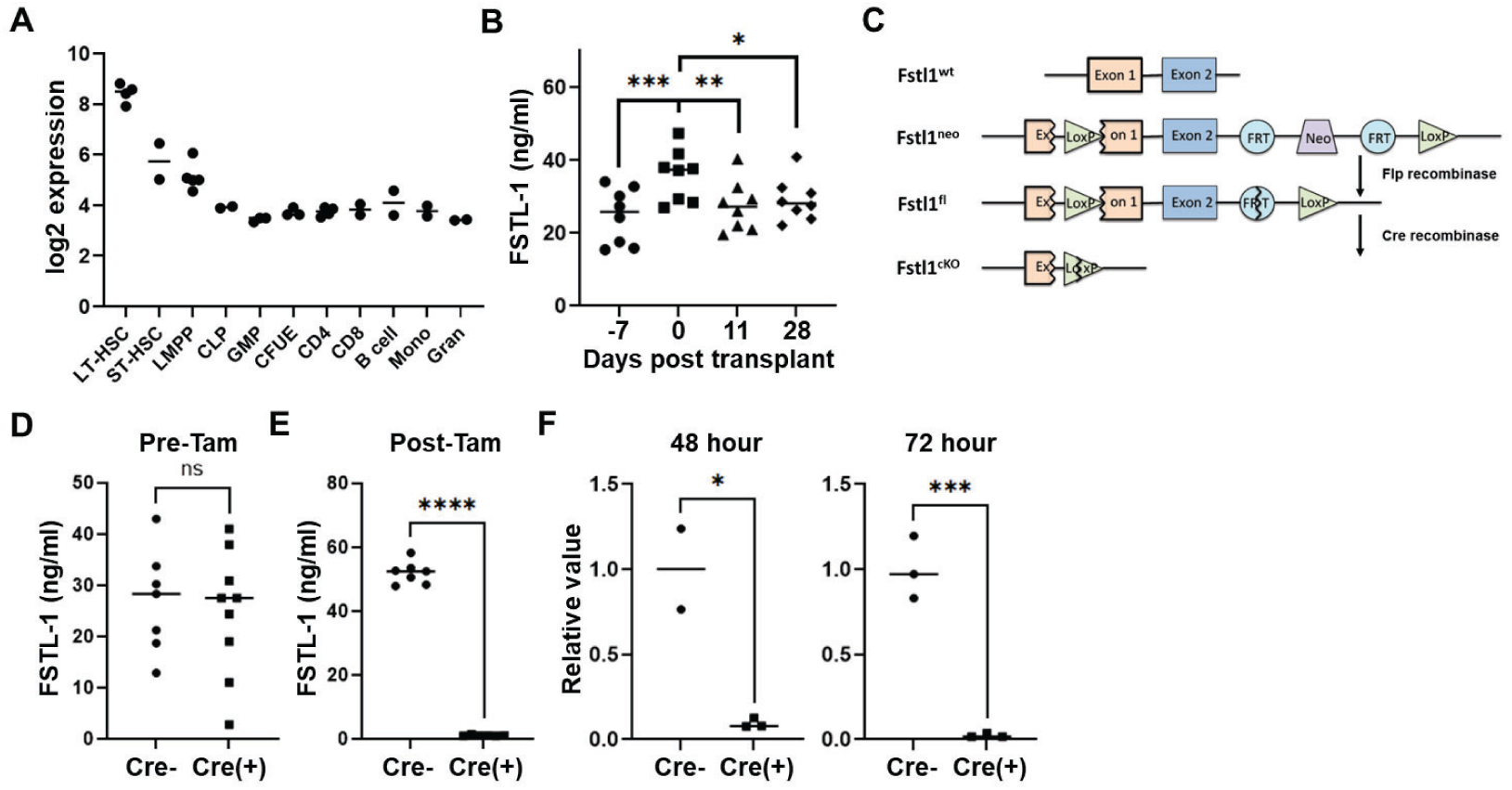
FSTL-1 in stem and stem progenitor cells. *A,* representative data on *FSTL1* expression in cells of the murine hematopoietic system (from BloodSpot.org). *B,* serum levels of FSTL-1 over the course of a syngeneic BM transplant. C, schematic outlining the strategy followed to insert LoxP sites into exon 1 and intron 2 of *Fstll.* Mice containing homozygously floxed *Fstll* alleles were subsequently bred to a strain expressing CreERT2 at the ROSA26 locus[34] to allow for conditional deletion of FSTL-1 in the presence of tamoxifen (FSTL-1 cKO). *D-E,* serum levels of FSTL-1 were measured in the mice (+/- CreERT2) prior to (D) or 7 days after **(E)** the first day of tamoxifen administration. *F,* Fstll^fl/fl^ mice (+/- CreERT2) were administered tamoxifen, followed 48 (left) or 72 (right) hours later by collection of whole BM cells and measurement of FSTL-1 levels by RT-PCR. Results are displayed relative to transcript levels in Cre negative control animals. *p<0.05, **p<0.01, ***p<0.001, ****p<0.0001

To investigate FSTL-1’s role in BM cells, we utilized a strain of mice bearing homozygous copies of loxP sites within exon 1 and downstream of exon 2 of the FSTL-1 gene. Fstl1^fl/fl^ mice were subsequently bred to the strain expressing CreERT2 under control of the Rosa26 locus [34], allowing for conditional deletion of FSTL-1 following tamoxifen exposure (FSTL-cKO). To test the efficacy of tamoxifen-driven, Cre-mediated deletion, mice were bled prior to tamoxifen administration and again on day 7 post-tamoxifen. Cre^neg^ mice given tamoxifen served as a vehicle control. Pre-tamoxifen, serum levels of FSTL-1 were identical between strains and independent of CreERT2 expression (Figure 1D). Following tamoxifen administration, serum FSTL-1 was essentially undetectable in Cre^pos^ (FSTL1-cKO) mice compared to Cre^neg^ littermate controls (Figure 1E). Evaluation of whole BM cells showed marked reduction of *Fstl1* transcript levels in FSTL1-cKO mice as early as 48 hours from the first dose of tamoxifen, with further decreases in *Fstl1* transcription as additional doses of tamoxifen were administered (Figure 1F). Together, these data demonstrate that 1) *Fstl1* is transcribed in whole BM cells, with the highest abundance in LT- and ST-HSCs, 2) serum levels of FSTL-1 change dynamically during the course of a standard BM transplant, 3) generation of a novel *Fstl1^fl^*^/fl^ strain allows for characterization of FSTL-1’s importance in specific populations of cells, and 4) breeding *Fstl1^fl^*^/fl^ mice to strains with ubiquitous expression of CreERT2 allows for conditional and significant deletion of *Fstl1* following tamoxifen exposure, including in BM cells.

### FSTL-1 deletion increases progenitor cell frequencies as well as numbers

Having constructed a method to achieve temporal deletion of *Fstl1*, we next determined the impact of FSTL-1 loss on BM progenitor cell number and frequency. We first focused our investigations on lineage negative, Sca-1+, c-kit+ (LSK) BM cells, given the suggested increase of *Fstl1* transcript levels within LT-and ST-HSC subpopulations (Figure 1A). The percentage of LSK+ cells in the BM of FSTL-1-cKO mice increased markedly 7 days after initial tamoxifen administration (Figure 2A-B), supported by a >2-fold increase in the total number of LSK+ progenitor cells recovered (Figure 2C). SLAM marker evaluation of LSK+ subpopulations demonstrated a marked increase in multipotent progenitor populations (MPP) MPP2 (CD48+CD150+) and MPP3/4 (CD48+CD150-) (Figure 2D-E), which was driven by enhanced expression of CD48, both in terms of percentage of CD48+ LSK cells (79.3% versus 92.7%, fl/fl versus FSTL1-cKO, respectively) as well as the median fluorescence intensity (MFI) of CD48 staining within the LSK gate (Supplemental Figure 2). As increased MPP percentages could arise from either an increased number of progenitor cells (i.e., more total MPP2 cells), or alternatively, from a decrease in the number of ST- and LT-HSCs, we measured the total number of each cell type present. Both MPP2 and MPP3/4 populations increased in total number in FSTL1-cKO mice, while LT-HSC numbers were equivalent between FSTL1-cKO mice and littermate controls, suggesting significant expansion of the committed progenitor pool. ST-HSC numbers, in contrast, decreased on day 7 following tamoxifen administration (Figure 2F), potentially as more cells transitioned into committed MPP2 and MPP3/4 progenitors. To determine if these day 7 results were preceded by changes in LT- and ST-HSC numbers at earlier time points, as might be expected if FSTL-1 were controlling HSC quiescence, we quantified stem and progenitor cell numbers at 48 hours, in mice having received two doses of tamoxifen. At this early 48-hour timepoint, there was a trend towards increased LT-HSC percentages in FSTL1-cKO mice, but no change in the percentage of any other stem or progenitor subpopulation (Supplemental Figure 3A). However, consistent with the day 7 data, the total number of LSK cells increased at 48 hours, which translated into elevated numbers of all HSPCs (Supplemental Figure 3B-C). Taken together, these results support a model where deletion of *Fstl1* disrupts HSPC homeostasis, marked first by an acute increase in the number of LT-HSCs (e.g., at 48 hours), with subsequent differentiation into more committed progenitor cells by day 7. LT-HSC levels return to baseline at this later timepoint, while ST-HSCs experience a requisite decrease as cells mature into more committed progenitor lineages.

**Figure 2:**
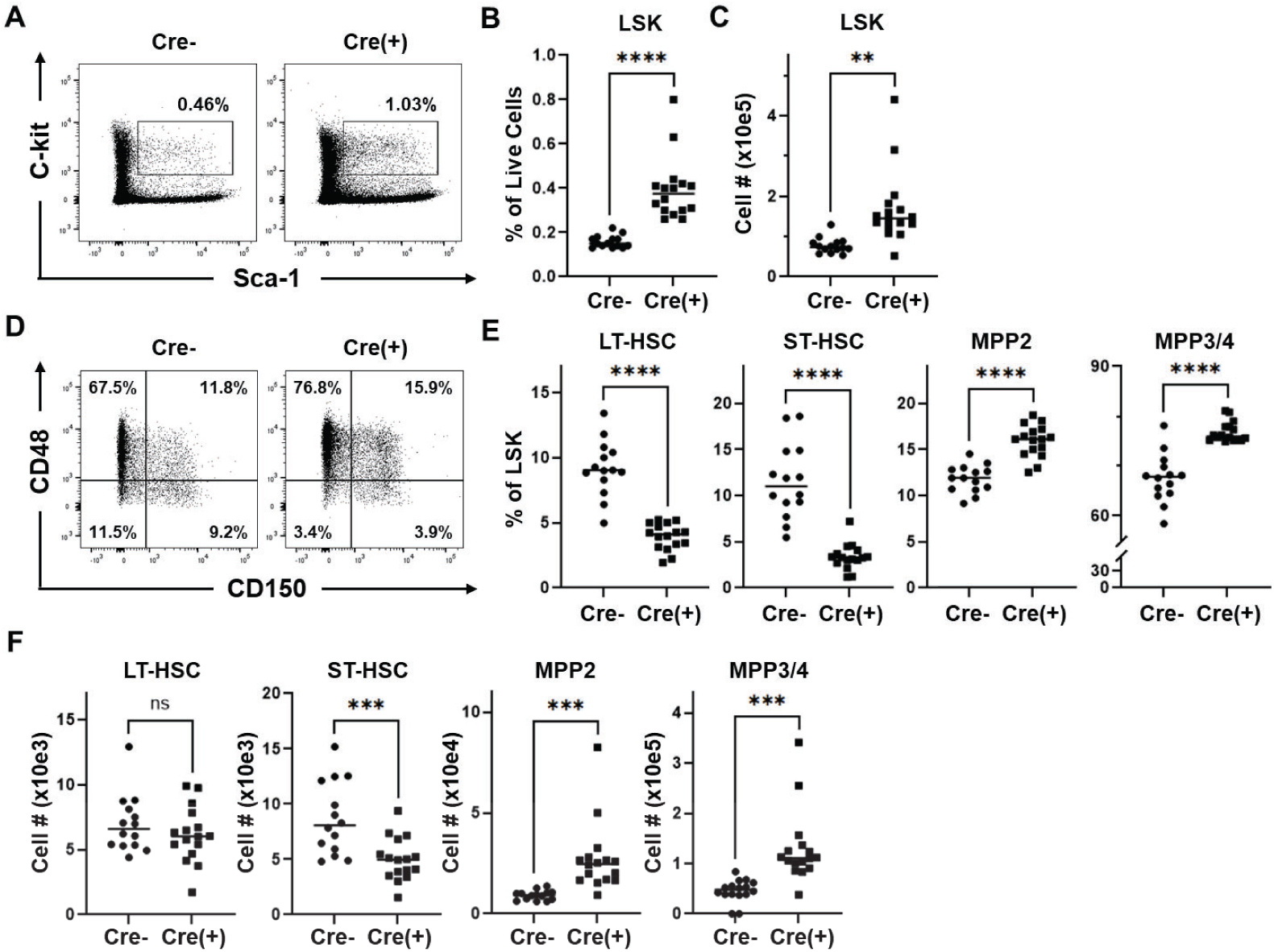
FSTL-1 deletion increases progenitor cell numbers and percentages. FSTL-1 cKO (Cre^pos^) and fl/fl (Cre^neg^) mice were treated with tamoxifen 4 times over 5 days, followed by analysis of BM stem and progenitor cells on day 7 post tamoxifen initiation. *A-C,* whole BM cells were stained with a lineage cocktail, and lineage negative cells were further gated on Sca-1+, c-kit+ cells **(A).** This LSK population was determined as a percentage of live cells in the BM (B) and as the total number of LSK cells (total number of viable BM cells multiplied by the percentage of LSK cells) **(C).** *D-F,* LSK cells were further gated using the SLAM markers CD48 and CD150 into subpopulations of MPP2 (CD48+CD150+), MPP3/4 (CD48+CD150-), ST-HSC (CD48-CD150-) and LT-HSC (CD48-CD150+). Representativestainingof fourteen to sixteen combined animals is shown in **(D),** while **(E)** displays values for individual mice. The total number of stem and progenitor cells for each subpopulation was calculated by multiplying their percentage against the total number viable BM cells recovered (F). n= 14 Cre^neg^ and 16 Cre^pos^ mice. **p<0.01, ***p<0.001, ****p<0.0001

### FSTL-1 ablation accelerates bone marrow reconstitution following sublethal irradiation

Having demonstrated both the presence of Fstl1 transcripts in early stem cell progenitors, as well as FSTL-1’s role in promoting BM homeostasis, we next sought to define the role played by FSTL-1 in recovery from BM injury. As outlined in Figure 3A, Fstl1^fl/fl^ mice (+/- Cre) were administered 4 doses of tamoxifen (over 5 days), followed by SLI with 450 cGy on day 14 following tamoxifen initiation. Two weeks post-irradiation, cells from the BM were recovered and the number and percentage of HSPCs analyzed. We first measured serum levels of FSTL-1. In Cre^neg^ mice, serum levels of FSTL-1 increased on day 1 post-irradiation and continued to be elevated until the time of BM collection on day +14 (Figure 3B). In contrast, serum levels of FSTL-1 were nearly undetectable in FSTL-1-cKO mice following tamoxifen administration, despite equivalent levels pre-tamoxifen. Consistent with an active role for FSTL-1 in suppressing BM differentiation following injury, the total number of BM cells recovered 14 days after irradiation increased markedly in FSTL-1-cKO mice (Figure 3C). Further evaluation revealed an additional increase in LSK cell percentages, which together with the increase in BM cell numbers, resulted in a >2-fold increase in total LSK cells recovered from the BM of FSTL-1-cKO mice (Figure 3D). There was no change in the percentage of specific HSPC populations as assessed by SLAM marker staining (Figures 3E-F), though the overall increase in LSK cells yielded a higher total number of all HSPC subpopulations, with elevations most marked in MPP2 and LT-HSC subpopulations, at >2.6- and 4.3-fold higher, respectively (Figure 3G). In total, these results demonstrate that serum levels of FSTL-1 are elevated following SLI and that absence of FSTL-1 following significant BM injury increases recovery of all HSPC cell types at early time points. Further, these imply that while all subpopulations of HSPCs benefit from an increase in BM cell number, LT-HSCs and MPP2 are most directly impacted by a lack of FSTL-1 in the recovering BM. Together, these findings imply that attenuation of FSTL-1-dependent pathways may serve as a beneficial means to increase and/or accelerate cell recovery following catastrophic BM injury, a concept that we envisioned could be leveraged therapeutically.

**Figure 3:**
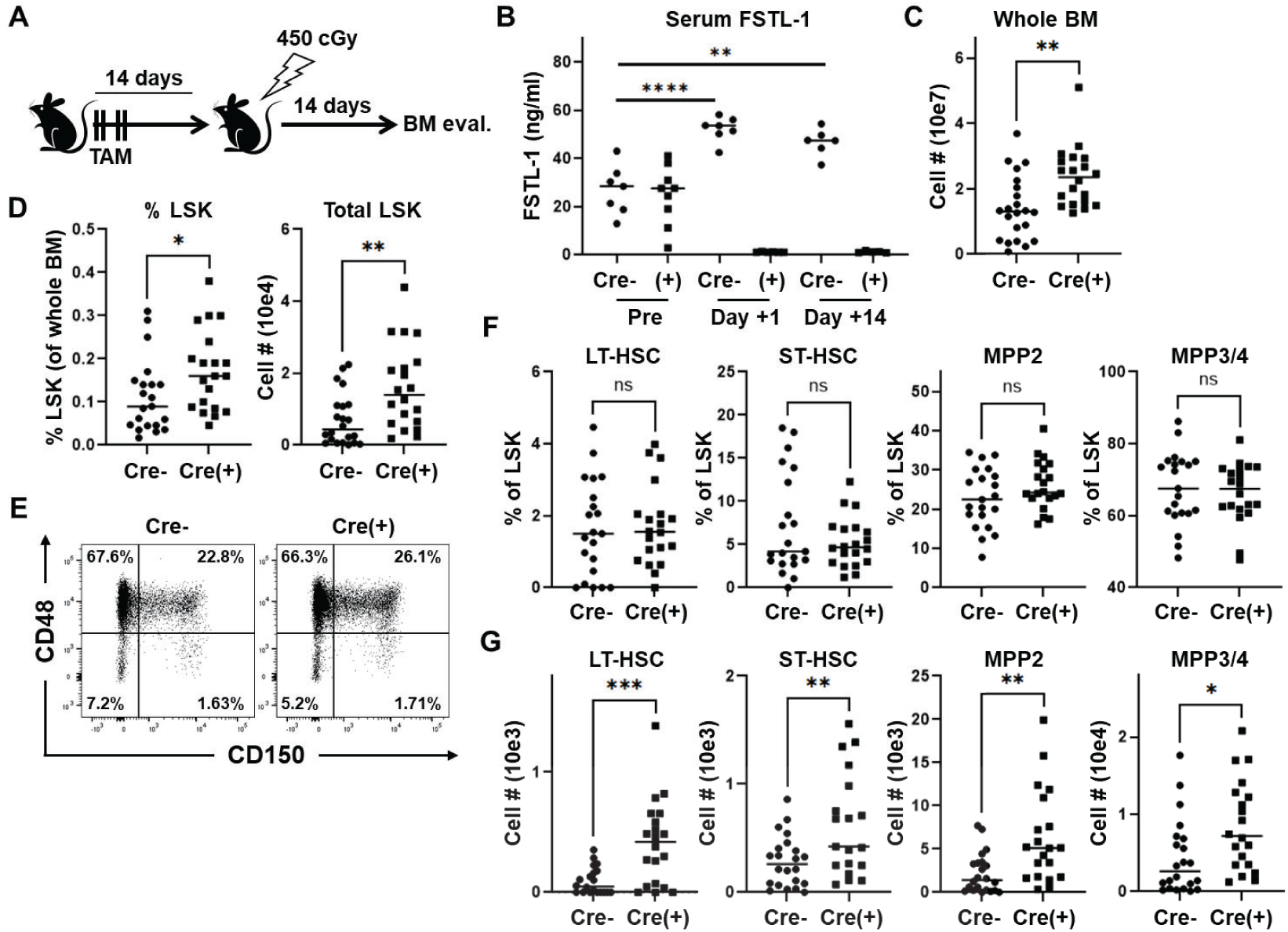
FSTL-1 ablation increases stem and progenitor cell numbers following sublethal irradiation. *A,* Fstll^fl/fl^mice (+/- Cre) were administered 4 days of tamoxifen, followed by 450 cGy of SLI 14 days later. Two weeks post-irradiation, BM cells were recovered and the number and percentage of HSPCs analyzed. B, serum levels of FSTL-1 were quantitated by ELISA either prior to tamoxifen administration (Pre) or on days +1 and +14 following SLI. *C-D,* the total number of BM cells were enumerated on day +14 post-irradiation from two femurs of the same mouse (C) followed by measuring the percentage (left) and total number (right) of LSK cells **(D).** *E-G,* Representative CD48CD150 SLAM staining is shown in (E), with dot plots displaying combined percentages from 22 Cre^neg^ and 20 Cre^pos^ animals. The stem and progenitor cell percentages for individual mice are plotted in (F). Total number of each subpopulation recovered was calculated as in Figure 2, where percentages of the whole were multiplied against the total number of viable BM cells recovered (G). n=7 Cre^neg^ and 8 Cre^pos^ mice in (B) and n=22 Cre^neg^ and 20 Cre^pos^ mice in the remaining plots. *p<0.05, **p<0.01, ***p<0.001, ****p<0.0001

### *In vitro* FSTL-1 blockade increases progenitor cells while decreasing short-term HSCs

To assess whether the improved BM recovery following FSTL-1 deficiency could be achieved therapeutically, we employed an orthogonal approach by inhibiting FSTL-1 using a polyclonal antibody generated against recombinant FSTL-1. We began by administering anti-FSTL-1 antibodies to overnight cultures of whole BM cells. Single cell whole BM suspensions were prepared from mouse femurs, washed in complete DMEM, and plated at 2e6 cells/well in in a 96-well round bottom plate. Cells were incubated overnight in the presence of rabbit IgG or rabbit polyclonal anti-sera specific to mouse FSTL-1 (#6942) at a final working concentration of 290ug/mL. Following overnight culture, cells were stained as described in Methods and flow cytometry performed the following day using our standard SLAM marker panel to differentiate HSCs and long-term progenitors (schematic outlined in Figure 4A). Overnight treatment with anti-FSTL-1 increased both the percentage and total number of LSK cells (Figure 4B). Within specific LSK subpopulations, FSTL-1 blockade decreased both LT-HSC and ST-HSC percentages, without impacting MPP2 or MPP3/4 frequencies (Figure 4C-D). These changes in cell percentages, combined with the increased total number of LSK cells, resulted in an overall decrease in total ST-HSCs, an increase in total MPP2 and MPP3/4 cells, and no change in LT-HSCs following 20 hours of FSTL-1 blockade (Figure 4E). Importantly, these results demonstrate that antibody blockade of FSTL-1, within HSPC populations, largely phenocopies the results obtained with genetic ablation of FSTL-1 (Figure 2F) and further suggests that antibody blockade of FSTL-1 could serve as a therapeutic modality to positively impact progenitor cell recovery following acute BM injury.

**Figure 4:**
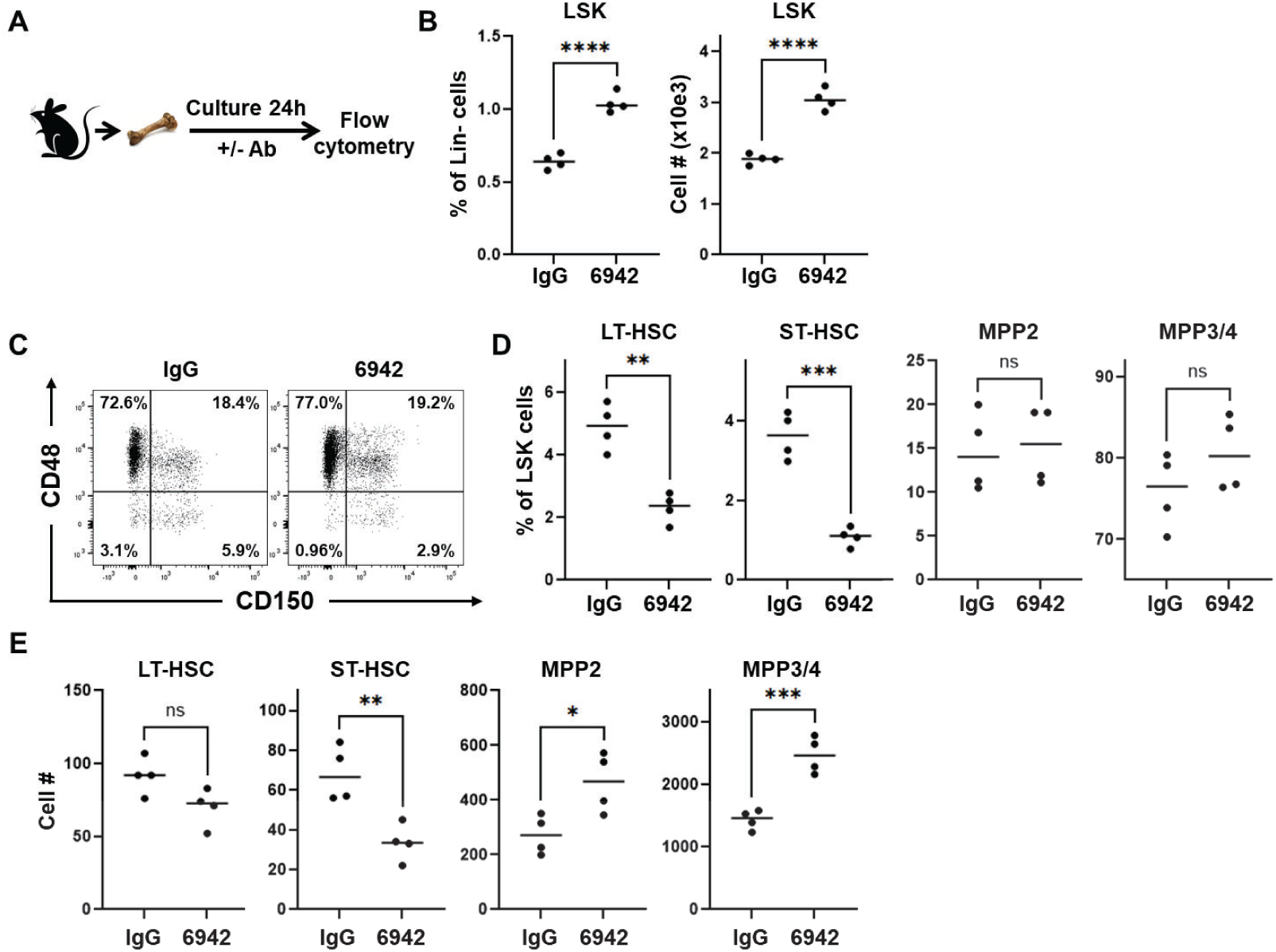
*In vitro* FSTL-1 blockade increases progenitor cell numbers while decreasing short-term HSCs. Bone marrow cells from WT C57BI/6 mice were cultured overnight in the presence of 6942, an FSTL-1 blocking antibody, or IgG control, with stem and progenitor cell percentages analyzed 24 hours later. *A*, experimental schematic. *B,* the percentage (left) and total number (right) of LSK cells at 24 hours of culture. C, representative flow plots of CD150 versus CD48 staining. *D-E,* The percentage **(D)** and total number **(E)** of cells in the HSC as well as MPP gates are shown. n=4 mice/group, with data combined from two independent experiments. *p<0.05, **p<0.01, ***p<0.001, ****p<0.0001

### *In vivo* blockade of FSTL-1 increases LT- and ST-HSC numbers following sublethal irradiation

Our initial data demonstrated that absence of FSTL-1 during the recovery from acute BM injury improved cellularity and increased the number of all HSC cell types (LT-HSC, ST-HSC, MPP2, and MPP3/4). To test whether *inhibiting* FSTL-1 in a clinically translatable manner (i.e. without genetic deletion) could enhance BM recovery, we attenuated FSTL-1 signaling via antibody blockade in our SLI model. Mice received 450 cGy of sublethal irradiation, followed by 6 subsequent doses of either control IgG or anti-FSTL-1 polyclonal antibody. Bone marrow cellularity, as well as HSPC subpopulations, were evaluated on day 14 post-irradiation (Figure 5A). Again, a strong trend towards increased BM cell numbers was observed in recipients of blocking anti-FSTL-1 antibody (Figure 5B), despite an equivalent percentage and total number of LSK cells (Figure 5C). Staining for CD48 and CD150 demonstrated an increased percentage of both LT- and ST-HSCs in anti-FSTL-1 treated mice (Figure 5D-E), which translated into greater total numbers of both LT- and ST-HSC cells. Surprisingly, this was not associated with changes in MPP2 of MPP3/4 on day 14 post-irradiation. Together, these data demonstrate that transient inhibition of FSTL-1, using a clinically translatable approach, increases both whole BM and HSC cell numbers at the critical time when mice are recovering from significant irradiation-induced BM injury, a result with potentially profound therapeutic implications.

**Figure 5:**
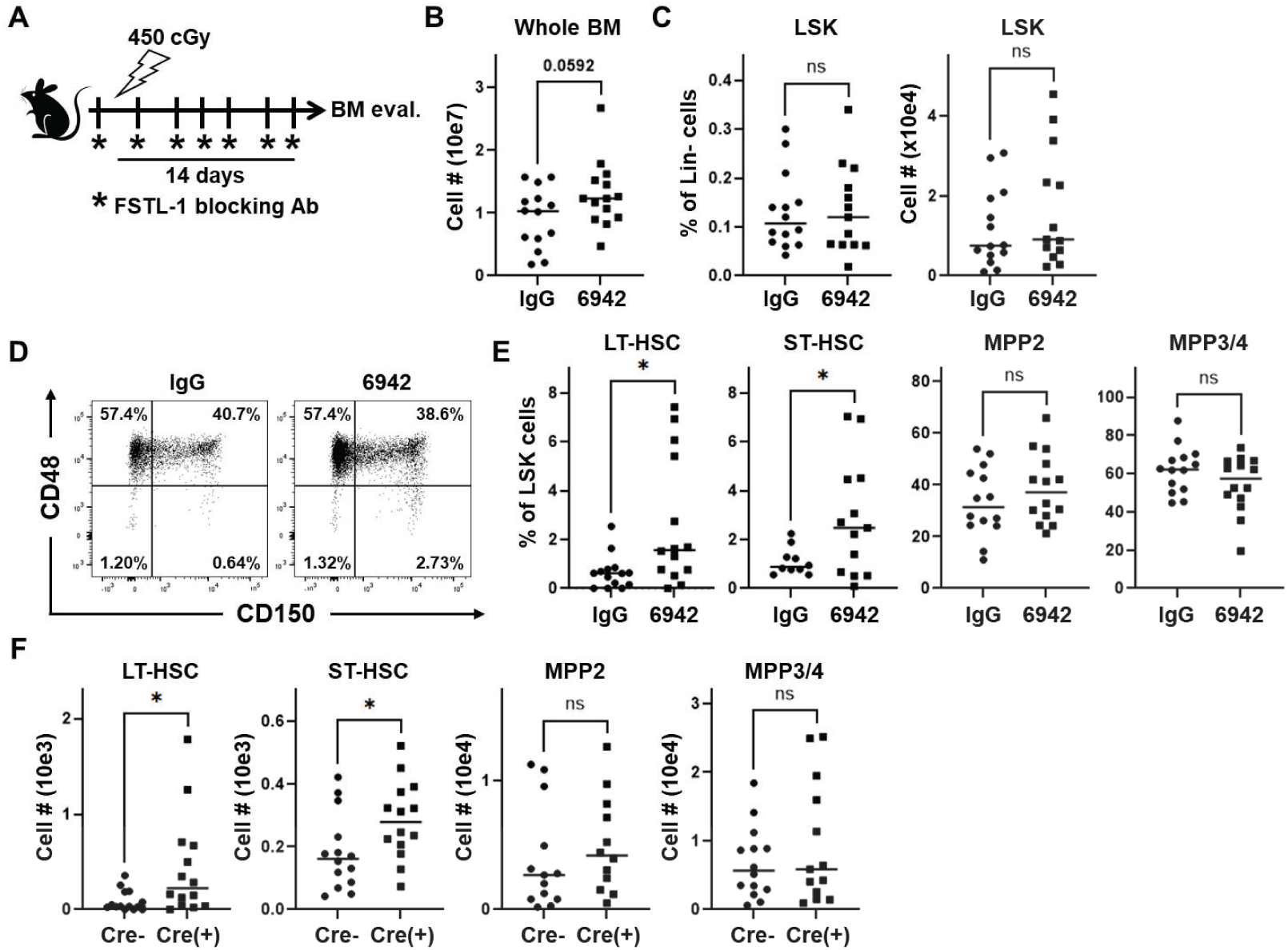
*In vivo* blockade of FSTL-1 increases LT- and ST-HSC numbers following sublethal irradiation. *A-C,* Wildtype C57BI/6 mice were sublet ha lly irradiated and then received anti-FSTL-1 blocking anti-sera three times per week as outlined in (A). Bone marrow cells were recovered from bilateral femurs and assayed for the total number of BM cells per animal (B) as well as the percentage (left) and total number (right) of LSK cells per mouse **(C).** *D-F,* Representative SLAM marker stainingfor one experiment is shown in (D), with the percentage of individual HSC and MPP subtypes displayed in (E), and the total number of HSC and MPP cells recovered from each animal demonstrated in (F). n= 16 mice/group, with data combined from two individual experiments. *p<0.05.

### Single cell RNA Sequencing of Lin^-^Sca1^+^Kit^+^ cells identifies a cell-specific FSTL-1 dependent transcriptomic program

Having identified a novel role for FSTL-1 in HSPC homeostasis and recovery from injury, we sought to explore potential FSTL-1-dependent mechanisms in HSPCs using single cell RNA Sequencing (scRNA Seq). Given that there was already a significant alteration in LSK and HSPC subpopulations by 7 days post-tamoxifen treatment (Figure 2B-C), we aimed to understand transcriptomic changes at an earlier time following *Fstl1* deletion. Forty-eight hours after Tamoxifen initiation, BM cells were collected from FSTL1-cKO and Cre^neg^ littermates for immediate sorting of LSK cells. 10,000 LSK cells per animal then underwent processing for single-cell RNA sequencing, as previously described [37]. Briefly, each sample was labeled using cell hashing antibodies, targeting recovery of ∼5,000 cells per sample. HTO-labeled cells were pooled and processed using the 10x Genomics Chromium 5ʹ Gene Expression Kit. Libraries were sequenced on an Illumina platform to a depth of approximately 50,000 reads per cell and reads were processed with Cell Ranger (v7.0) and analyzed in Seurat (v5.2.0).

Using gene signatures, UMAP analysis identified 8 distinct clusters within LSKs (Figure 6A) designated as: HSCs, Proliferating cells, Neutrophils, Megakaryocytes, Erythroid cells, Basophils, Lymphocytes, and Monocytes; a representative heatmap is shown in Figure 6B. Having annotated discrete LSK clusters, we next sought to discern differences in cluster representation along the LSK cell trajectory as a function of FSTL1 deficiency. Principle component analysis (PCA) identified a four-armed pseudotime map, wherein FSTL1-cKO (Cre^pos^) mice were relatively enriched in overall cellularity, most notably within the HSC and Proliferating cell clusters (Figure 6C). Differential expression of *Fstl1* was most abundantly identified within the HSC (adjusted p-value 1.30e-14) and Proliferating cell (adjusted p-value 3.28e-14) clusters, as well as within Megakaryocytes (adjusted p-value 8.67e-6). Fidelity of this original finding, as well as validation of Fstl1 elimination, was confirmed by the significant reduction of *Fstl1* expression in these clusters in Cre^pos^ (FSTL1-cKO), but not in Cre^neg^, mice (Fig 6D). Comparing differentially expressed genes (DEG) between WT and FSTL1-cKO LSKs identified 429 significantly altered transcripts across LSK clusters, with the greatest numbers concentrated in the HSC (178 total, 112 downregulated, 66 upregulated), Neutrophil (148 total, 110 downregulated, 38 upregulated), and Proliferating cell clusters (90 total, 55 down, 35 upregulated) (Supplemental Table I).

**Figure 6:**
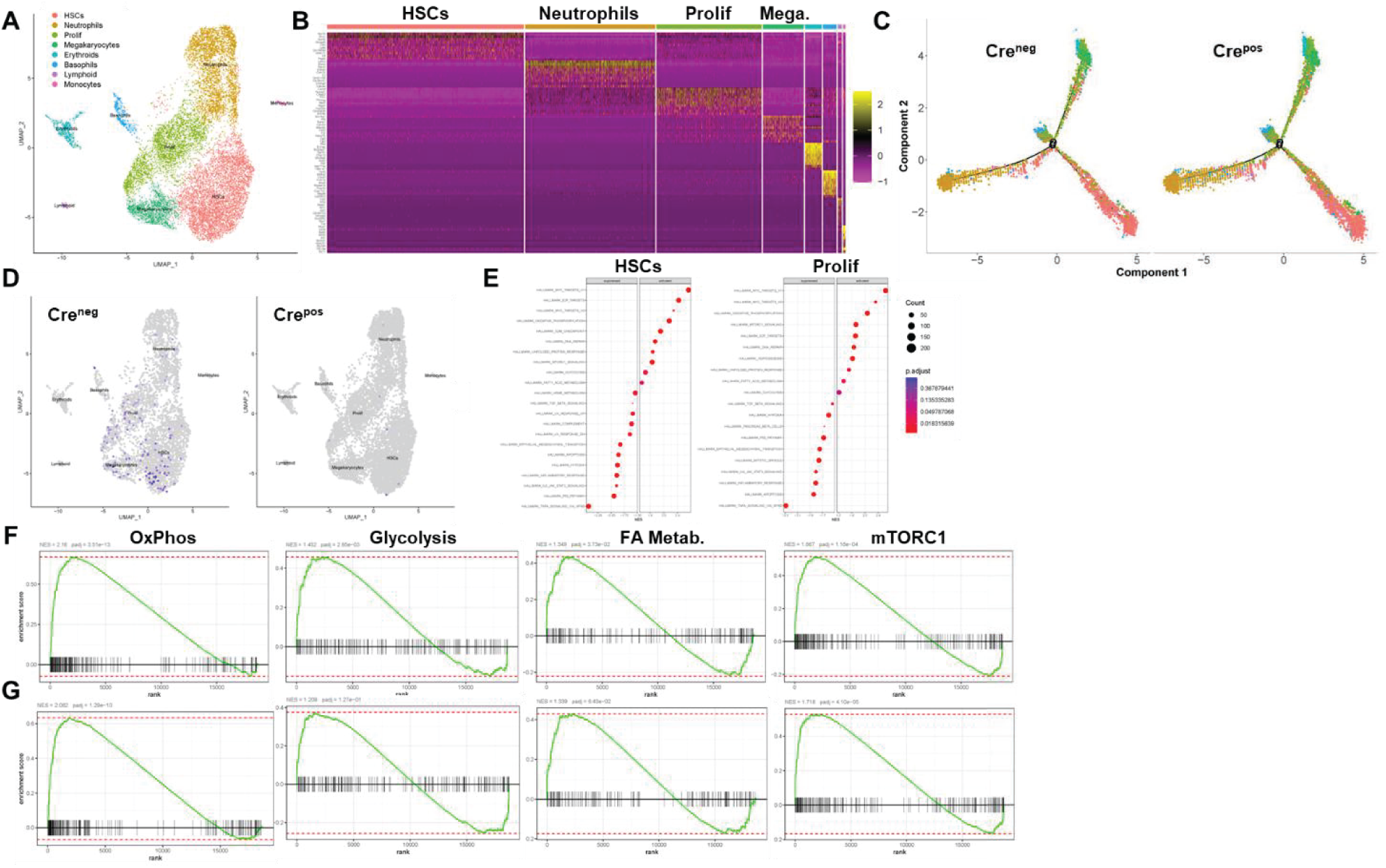
Single cell RNA sequencing identifies a role for *Fstll* in regulation of cell cycle progression and metabolism in specific HSPC populations. FstI^fl/fl^ mice (+/- Cre) were administered tamoxifen for two consecutive days and BM harvested 48hrs afterthe first dose (n=2/genotype). BM cells underwent staining to identify the LSK populations for sorting by FACS. 10,000 FACS sorted LSK cells from Cre^neg^ and Cre^pos^ mice were then processed for single-cell RNA sequencing as outlined in Methods. *A-D,* UMAP analysis identified 8 individual clusters as designated in (A). Heatmap (B) il I ust rates the differential expression of cluster-defining transcripts. Trajectory clusters in Cre^neg^(WT) and Cre^pos^ (FSTL1 cKO) LSK cells identified enhanced representation of HSC and Prolife rati ng cel Is populationsin Cre^pos^ mice **(C).** Comparing *Fstll* expression in LSK cells from in Cre^neg^ (WT) and Cre^pos^(FSTL1 cKO) mice by scRNA-seq analysis (D). Gene set enrichment analysis (GSEA) showing the top 10 most-suppressed and most-activated pathways for HSCs (left) and Proliferating cells (right) **(E).** *F-G,* GSEA-derived ribbon plots for selected cell cycling and metabolic pathways for HSCs (F) and Proliferating cells **(G).** NES = Normalized Enrichment Scores; padj. = adjusted p-value.

Recognizing that HSC and Proliferating cell clusters both experienced the biggest reduction in *Fstl1* expression, along with the majority of transcriptional changes, we hypothesized that these transcriptional changes were likely to be FSTL-1-dependent. In striking similarity, cells lacking *Fstl1* from both HSC and Proliferating cell clusters significantly increased expression of transcripts for *Zfp36*, *Jun*, *Fos*, *Nr4a1, Klf6,* and *Ier2*, while similarly decreasing representation of both *Mif* and *Ranbp1* (Supplemental Tables II and III). To elucidate FSTL-1-dependent *<u>pathways</u>*, gene set enrichment analysis (GSEA) was performed. Within both the HSC and Proliferating cell clusters, pathways most upregulated related to proliferation (Myc targets, EF2 targets, G2M checkpoint) and cellular metabolism (oxidative phosphorylation (OxPhos), mTORC1 signaling, glycolysis, fatty acid metabolism). In contrast, pathways related to inflammation and apoptosis (TNFa signaling via NF-kB, IL6-JAK-STAT3 signaling, inflammatory response) were significantly suppressed (Figure 6E). Together these data suggest that cells within the LSK compartment, which lack FSTL-1, reduce inflammatory and apoptotic pathways, while simultaneously increasing proliferation and pathways associated with cellular metabolism.

To support this conclusion, we further analyzed enrichment of metabolism related pathways specifically within the HSC cluster, identifying significant Normalized Enrichment Scores (NES) in FSTL-cKO cells for OxPhos (NES 2.16, p-adj 3.51e-13), glycolysis (NES 1.452, p-adj 2.85e-3), fatty acid metabolism (NES 1.349, p-adj 3.73e-2), and mTORC1 signaling (NES 1.667, p-adj 1.16e-4) (Figure 6F). Analysis of the Proliferating cell cluster revealed a very similar NES increase for OxPhos (NES 2.062, p-adj 1.29e-10), fatty acid metabolism (NES 1.339, p-adj 6.43e-2), and mTORC1 signaling (NES 1.718, p-adj 4.10e-5) (Figure 6G). Thus, homeostatic deletion of FSTL-1 specifically alters transcriptional signatures related to cellular proliferation and metabolism within HSC and Proliferating cell populations.

Cumulatively, these studies identify select expression of FSTL-1 within specific HSPC subtypes. Further, inhibition of FSTL-1’s actions, through either genetic deletion or anti-FSTL-1 antibody-treatment, impacts the balance between long-term repopulating cells and committed precursor populations, a phenomenon apparent during homeostasis that is further exaggerated following BM injury. Finally, unbiased transcriptomic analysis demonstrated FSTL-1-mediated impacts on HSCs and proliferating subtypes of HSPCs, including accentuation of proliferation and metabolism-associated pathways, with concurrent suppression of inflammatory signaling. Altogether, these results suggest a prominent role for FSTL-1 in regulating cell-specific HSPC function, highlighting a mechanism amenable to therapeutic intervention.

## Discussion

A short hairpin RNA screening strategy, delivered to repopulating HSPCs, first implicated a potential role for FSTL-1 in HSC biology [35]. In that study, knockdown of Fstl1 decreased *in vivo* HSPC repopulating ability and diminished colony-forming potential, particularly within the CFU-GM/G/M and CFU-GEMM populations. However, no further characterization was provided. Later, a similar role for FSTL-1 was suggested, downstream of the chromatin remodeler Znhit1 [38]. In that study, Znhit1 loss resulted in acute expansion of HSCs and simultaneous decrease in *Fstl1* expression, alongside decreased expression of HSC quiescence genes including *Foxo1, Pten, Nr4a1*, and *Klf4*. The current study extends this research by taking a more direct approach, specifically suppressing Fstl1 expression at defined times, and thereby allowing a specific evaluation of FSTL-1-related changes in HSC populations during both homeostasis and in response to sublethal BM injury. This approach avoids the transient and somewhat inconsistent inhibition seen with shRNA, while also circumventing the potential confounding effects of impacting multiple targets, as seen with Znhit1 deletion and associated chromatin remodeling. Notably, all three studies consistently support the concept of FSTL-1 playing a vital role in modulating HSC quiescence and repopulation potential.

HSCs have been defined on the basis of two essential properties, self-renewal and the capacity for multipotent differentiation [12]. This definition distinguishes HSCs from multipotent progenitor cells (MPPs), with HSCs having long-term renewal capacity, while MPPs exhibit lineage specific output and reduced self-renewal [39]. In the classic model, long-term (LT)-HSCs differentiate into short-term (ST)-HSCs, which subsequently differentiate into subgroups of MPPs; which demonstrate only short-term repopulation capacity. A large body of work indicates that differentiation of HSCs into multipotent progenitors (MPPs) is a tightly controlled, stepwise process governed by both intrinsic transcriptional programs and extrinsic signals delivered by the BM niche. Steps in this process include appropriate metabolic activation, controlled cell-cycle entry, and supportive stromal and endothelial interactions [40]. While this classic interpretation frames the elegance of HSPC differentiation, it likely oversimplifies the complexity of HSPC biology, with recent single-cell and multi-omics studies revealing substantial heterogeneity within the HSC-to-MPP transition and distinct subpopulations primed for myeloid or lymphoid differentiation even before full lineage commitment [41]. Further, there is evidence that acquisition of lineage fates might actually be a continuous process [42] and that MPPs, rather than HSCs, are the main drivers of hematopoiesis in adulthood [2], while other work even argues against the existence of strict boundaries between stem and progenitor cells [43]. Collectively, the current and historical evidence frames the HSC-to-MPP differentiation as a plastic yet vulnerable transition state, influenced by the integration of signaling, metabolic, and microenvironmental cues. Our data support a role for FSTL-1 in maintaining cells in the LT- and ST-HSC states, with inhibition of FSTL-1 promoting differentiation into more specialized MPP cells (Figures 2E and 4D,E). Furthermore, in contrast to G-CSF, or CXCR4 antagonists like plerixafor, which impact a heterogeneous population of hematopoietic progenitors (of which LT-HSCs represent only a minor component) [44], the impact of FSTL-1 amelioration seems to be much more specific to HSCs, based on both our differentiation analysis and the pattern of *Fstl1* cellular expression in our scRNASeq studies (Figures 1A and 6D).

HSCs exhibit a dynamic yet highly regulated metabolic program, which is tightly coupled to their quiescence, self-renewal, and differentiation potential. Recent studies emphasize *metabolic plasticity* as a central determinant of stem cell fate [45–47], where in the ambient hypoxia of the BM niche, quiescent HSCs primarily rely on anaerobic glycolysis, with minimal contributions from mitochondrial OxPhos and thus limited reactive oxygen species generation [13]. Upon activation or differentiation, stem cells shift toward increased mitochondrial respiration and elevated OxPhos, to meet a surge in bioenergetic demands and biosynthetic requirements [48]. In this context, amino acid and fatty acid metabolism contribute to acetyl-CoA production, which in turn regulates HSC asymmetric division and subsequent lineage commitment [49, 50]. This previous research underscores the fact that HSC metabolism is not static, but represents a malleable process integrating glycolytic, mitochondrial, and accessory metabolic pathways to balance HSC maintenance, self-renewal, and differentiation [51, 52]. Our findings agree with these reports, positing a role for FSTL-1 in promoting HSC quiescence, while also demonstrating that absence of FSTL-1 leads increases HSC proliferation and differentiation, concurrent with transcriptomic changes in both glycolysis- and OxPhos-related genes.

FSTL-1 has been well studied for its potential to support stem cell function and heal damaged tissues in other contexts [33, 53]. In heart disease models, protein delivery of human FSTL-1 via an epicardial patch stimulated proliferation of pre-existing cardiomyocytes (CMs) [25] and helped regenerate ischemic heart tissue [54]. This approach also promoted, in a hypoxia injury model, the *in vitro* proliferation of CMs matured from human induced pluripotent stem cells [55]. In glioblastoma, glioma stem cells utilize autocrine FSTL-1 to maintain activation of PI3K-AKT signaling, a step critical in maintenance of their self-renewal capacity [56]. A similar pattern is seen in breast cancer cells, where stemness is maintained through an FSTL-1/integrin Beta3/Wnt signaling pathway [27]. In the spleen, lineage development of a unique MPP population is regulated by NR4A1 [57], a transcription factor that also restrains HSC proliferation and whose expression and function are regulated by FSTL-1 during immune cell homeostasis in the lung [28]. Additionally, FSTL-1 enhances the therapeutic efficiency of mesenchymal stem cells (MSCs), both in treating myocardial infarction [26] but also during liver fibrotic therapy, where MSCs provide an immunosuppressive effect on inflammatory macrophage responses [58]. While FSTL-1 has been implicated in various contexts of tissue repair and stem cell homeostasis, the present study on FSTL-1 aligns most closely with those of the glioblastoma and breast cancer stem cells, where basal levels of FSTL-1 work in an autocrine fashion in LT-HSCs to promote a program of quiescence and self-renewal, while disruption of FSTL-1 promotes LT-HSC differentiation transiently into ST-HSCs, then into more committed MPPs less capable of self-renewal.

We are not the first group to demonstrate a therapeutic benefit of targeting FSTL-1. In liver fibrosis, inhibition of FSTL-1 via blocking antibody both attenuated liver fibrosis and delayed hepatic stellate cell activation. These changes occurred via a decrease in TGF-beta signaling, a result consistent with the suppression of TGF-beta signaling seen by scRNASeq in FSTL-1 KO HSPCs (Fig 6E). In pulmonary fibrosis, FSTL-1 mediates the myofibroblast pro-fibrotic response, and antibody inhibition of FSTL-1, reduced fibrosis *in vivo* [24]. Our results are consistent with anti-FSTL-1 treatment promoting HSC differentiation and subsequent accumulation of MPPs, observed in both the FSTL-1 genetic deletion and antibody models (e.g., compare Figure 2F and 4E). Further, we anticipate that FSTL-1 blockade may offer distinct therapeutic advantages over currently approved therapies. G-CSF speeds recovery of specific myeloid lineage precursors while plerixafor promotes egress of lineage differentiated cells from the BM. These approaches, however, do little to facilitate the differentiation or proliferation of early-stage HSPCs, a critical void which could be met by FSTL-1-targeted therapies.

Therapeutically, increasing BM output would be expected to benefit both recovery from acute BM insult (e.g., radiation exposure) as well as promoting accelerated BM recovery in situations of iatrogenic BM injury (i.e., conditioning from blood and marrow transplantation). Furthermore, inflammatory cytokines are known to work directly on HSPCs to promote ‘emergency hematopoiesis’ during serious infections [59], a process which may be further amplified by FSTL-1 blockade. Finally, inhibition of FSTL-1 could be envisioned to treat the subset of sepsis patients who experience profound BM suppression following an episode of sepsis. This is particularly important as G-CSF, despite attenuating inflammatory responses, has shown little efficacy in improving clinical outcomes [60, 61], even though HSPC dysfunction was not present in these patients [62].

## Conclusions

In summary, this study highlights a novel role for FSTL-1 in controlling the homeostatic and differentiation potential of HSPCs. Specifically, elimination of FSTL-1 facilitates an increase in BM cellularity under homeostatic conditions, an influence that increases following the significant injury to the BM wrought by substantial irradiation. Importantly, these changes are similar following treatment with an FSTL-1 blocking antibody. Finally, scRNASeq demonstrates that *Fstl1* expression is limited to distinct cellular subsets within the HSPC compartment, and with a notable change in metabolic pathways in these same cells in the absence of FSTL-1. In total, these results not only deepen our fundamental knowledge of FSTL-1 biology, but could help to predict ways in which to influence HSC and HSPC biology to therapeutic advantage.

## List of abbreviations

BM: Bone Marrow
CLR: Centered Log Ratio
Cre: Cre Recombinase
DEG: Differentially Expressed Gene
DMEM: Dulbecco’s Modified Eagle Media
ERT2: Estrogen Receptor T2
ELISA: Enzyme-linked Immunosorbent Assay
FBS: Fetal Bovine Serum
FSTL-1: Follistatin-like 1
GSEA: Gene Set Enrichment Analysis
HEPES: 4-(2-hydroxyethyl)-1-piperazineethanesulfonic acid
Hprt1: Hypoxanthine phosphoribosyltransferase 1
HSC: Hematopoietic Stem Cell
HSPC: Hematopoietic Stem and Progenitor Cell
HTO: Hashtag Oligo
IACUC: Institutional Animal Care and Use Committee
IgG: Immunoglobulin G
i.p.: Intraperitoneal
KLF4: Krüppel-like Factor 4
LSK: Lineage negative, Sca-1+c-Kit+
LT-HSC: Long-term Hematopoietic Stem Cell
MAST: Model-based Analysis of Single-cell Transcriptomics
MFI: Median Fluorescence Intensity
MPP: Multipotent Progenitor
mTORC1: Mammalian Target of Rapamycin Complex 1
NES: Normalized Enrichment Score
NR4A1: Nuclear Receptor subfamily 4 group A member 1
OxPhos: Oxidative Phosphorylation
Padj: Adjusted p value
PBS: Phosphate-buffered Saline
PCA: Principal Component Analysis
RNA: Ribonucleic Acid
RT-PCR: Reverse Transcriptase Polymerase Chain Reaction
scRNASeq: Single cell RNA Sequencing
SLI: Sublethal Irradiation
SNN: Shared Nearest Neighbor
ST-HSC: Short-term Hematopoietic Stem Cell
TGF-β: Transforming Growth Factor – Beta
TMB: 3,3’,5,5’-tetramethylbenzidine
UMAP: Uniform Manifold Approximation and Projection

## Declarations

### Ethics approval and consent to participate

All mice were housed in a University of Pittsburgh pathogen-free facility in accordance with the Institutional Animal Care and Use Committee (IACUC) established guidelines. Further, all experiments were performed under University of Pittsburgh IACUC approved protocols.

### Consent for publication

Not applicable.

### Availability of data and materials

The scRNASeq datasets generated during the current study will be made available in the GEO digital repository, under accession number [To be filled in upon acceptance of the manuscript]. Data from the BloodSpot dataset can be found at Bloodspot.org using the query “‘mouse normal hematopoietic system”. It is also referenced here [36].

### Author’s contributions

CAB designed and performed experiments, analyzed data, interpreted all experiments, and wrote and edited the paper with BTC. MH designed and performed experiments, analyzed data, and edited the manuscript. NS performed experiments and edited the manuscript. DL analyzed data. KC designed and performed experiments, supervised data analysis, and edited the manuscript; BTC designed and supervised experiments, analyzed and interpreted all data, and wrote and edited the manuscript with CAB. All authors read and approved the final manuscript.

## Acknowledgements

Not applicable.

