## Supplemental Figures and Tables for "A Novel Role for FSTL-1 in Hematopoietic Stem Cell Homeostasis and Post-injury Regeneration"

Supplemental Figures: 3

### Supplemental Figures

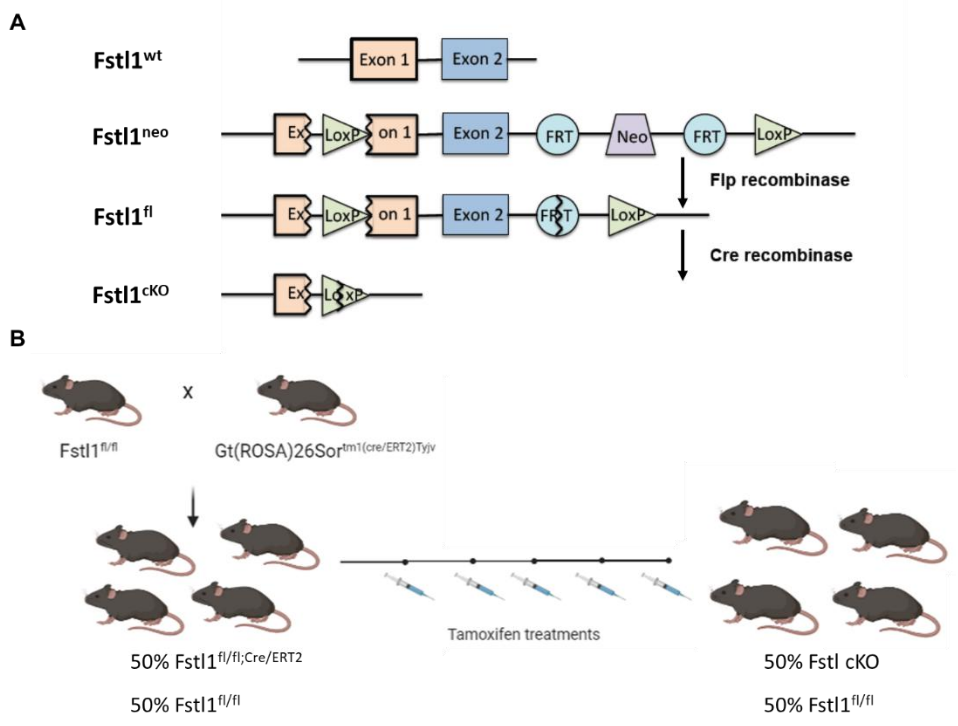

**Supplemental Figure 1 – Generation of FSTL-1 deficient mice.** *A,* Floxed *Fstl1* generation is depicted schematically, and has been described previously [1]. *B, Fstl1$^{fl/fl}$* mice were then bred with Gt(ROSA)26Sor$^{tm1(Cre/ERT2)Tyjv}$ to generate FSTL1$^{fl/WT;\ Cre/ERT2}$ mice; these mice were bred back to *Fstl1$^{fl/fl}$* mice to generate *Fstl1$^{fl/fl;\ Cre/ERT2}$*. Thereafter *Fstl1$^{fl/fl;\ Cre/ERT2}$* and *Fstl1$^{fl/fl}$* mice were intercrossed to generate litters containing ~50% *Fstl1$^{fl/fl;\ Cre/ERT2}$* and 50% *Fstl1$^{fl/fl}$* mice. To delete the *Fstl1* allele, Cre-recombinase was activated by treating mice with 4mg tamoxifen in corn oil via intraperitoneal (i.p.) injection for two consecutive days, were give a day of rest, followed by two more consecutive days of 4 mg i.p. tamoxifen administration. This approach yielded ~50% *Fstl1* conditional knockout (cKO) mice (lacking FSTL-1) and ~50% FSTL-1 sufficient (*Fstl1$^{fl/fl}$*) littermate controls.

.

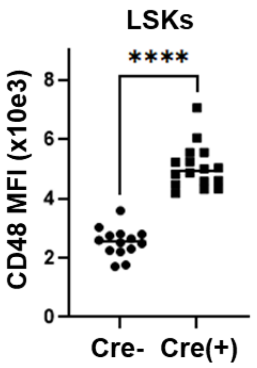

**Supplemental Figure 2 – CD48 levels increase in LSK cells from FSTL-1 deficient mice.** CreERT2$^{pos}$ and Cre$^{neg}$ animals were administered tamoxifen for 4 out of 5 days. Seven days from initial tamoxifen administration, whole BM cells were recovered, stained for extracellular markers, and gated on Lin$^{neg}$Sca-1+c-kit+ (LSK) cells. This process was followed by assessment of CD48 levels on LSKS via flow cytometry and determination of median fluorescence intensity (MFI) ****p<0.0001.

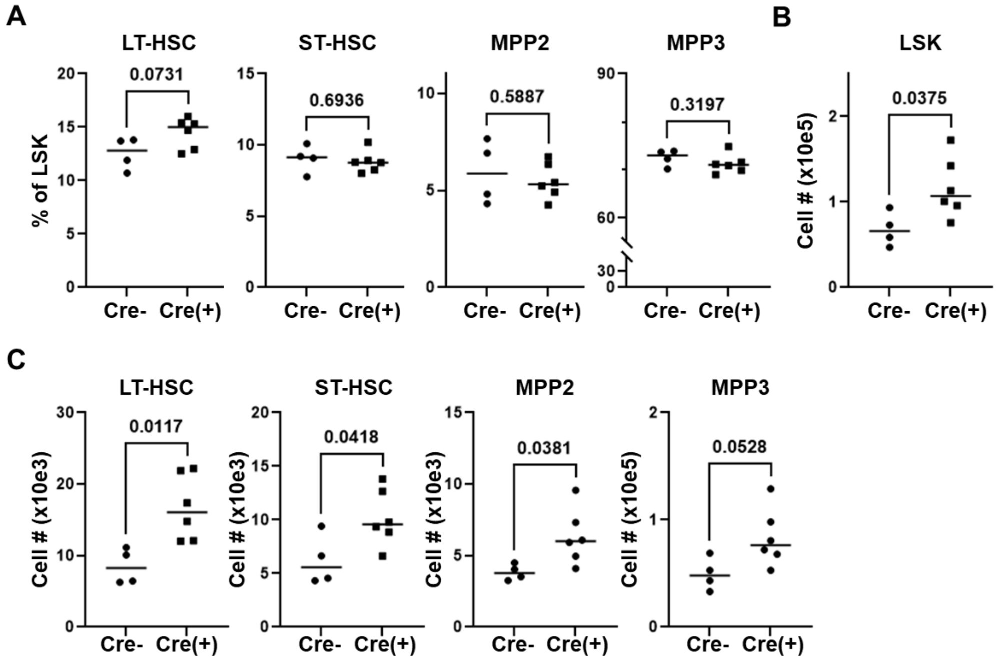

**Supplemental Figure 3 – Stem and progenitor cell numbers increase within 48 hours of FSTL-1 deletion.** *A-C*, CreERT2$^{pos}$ and Cre$^{neg}$ animals were administered tamoxifen over two days. Forty-eight hours from the first dose of tamoxifen, whole BM cells were recovered, counted, and stained. These steps were followed by gating on Lin$^{neg}$Sca-1+c-kit+ (LSK) cells, with further assessment of stem and progenitor cell subtypes (**A**). Total cell numbers were then determined by multiplying the percentage of each cell subpopulation by the total number of viable BM cells recovered, shown for both LSK cells (**B**) and for each of the stem and progenitor cell subpopulations (**C**).

### Supplemental Table I: Differentially Expressed Genes

| Cluster | DEGs (n) | Up (in FSTL1 Cre+) | Down (in FSTL1 Cre+) | Cre$^{neg}$ *Fstl1* expression | Cre$^{pos}$ *Fstl1* expression | padj (Fstl1) |
|---|---|---|---|---|---|---|
| Basophils | 1 | 1 | 0 | 0.022 | 0.004 | 1 |
| Eryhtroids | 3 | 2 | 1 | 0 | 0 | 1 |
| Lymphoids | 0 | 0 | 0 | 0 | 0 | 1 |
| Monocytes | 0 | 0 | 0 | 0 | 0 | 1 |
| Neutrophils | 148 | 38 | 110 | 0.012 | 0 | 0.2880337 |
| Megakaryocytes | 9 | 6 | 3 | 0.056 | 0 | 8.67E-06 |
| Proliferating | 90 | 35 | 55 | 0.052 | 0.001 | 3.28E-14 |
| HSC | 178 | 66 | 112 | 0.03 | 0.001 | 1.30E-14 |
| **Total DEGs** | **429** | | | | | |

**Supplemental Table II - DEGs from HSC cluster**

| | p_val | avg_log2FC | pct.1 | pct.2 | p_val_adj |
|---|---|---|---|---|---|
| Zfp36 | 2.21E-76 | -0.667569835 | 0.772 | 0.907 | 4.14E-72 |
| Jun | 2.22E-71 | -0.791212811 | 0.656 | 0.801 | 4.16E-67 |
| Fos | 3.10E-41 | -0.693168138 | 0.632 | 0.767 | 5.81E-37 |
| Junb | 7.08E-37 | -0.503322787 | 0.698 | 0.79 | 1.33E-32 |
| Mif | 5.18E-36 | 0.310469159 | 0.974 | 0.941 | 9.71E-32 |
| Pde4b | 1.12E-34 | -0.429265926 | 0.695 | 0.784 | 2.09E-30 |
| Ctla2a | 1.39E-33 | -0.473989776 | 0.842 | 0.907 | 2.61E-29 |
| Ier2 | 8.96E-33 | -0.434914189 | 0.726 | 0.808 | 1.68E-28 |
| Dusp1 | 2.18E-31 | -0.546628482 | 0.357 | 0.516 | 4.08E-27 |
| Klf6 | 8.70E-27 | -0.451125038 | 0.13 | 0.244 | 1.63E-22 |
| Ccl4 | 5.65E-26 | -0.638006568 | 0.187 | 0.307 | 1.06E-21 |
| Btg2 | 2.41E-25 | -0.432309171 | 0.265 | 0.404 | 4.51E-21 |
| Nop10 | 1.26E-24 | 0.253072255 | 0.93 | 0.854 | 2.36E-20 |
| Eif5a | 3.15E-24 | 0.196562653 | 0.987 | 0.975 | 5.89E-20 |
| Ppia | 1.21E-22 | 0.139426199 | 0.999 | 0.999 | 2.27E-18 |
| mt-Co1 | 6.20E-22 | -0.177417267 | 0.999 | 0.999 | 1.16E-17 |
| Rgs1 | 8.91E-22 | -0.505231261 | 0.303 | 0.424 | 1.67E-17 |
| 2410006H1 | 4.03E-20 | -0.298765888 | 0.894 | 0.92 | 7.55E-16 |
| Ran | 5.73E-19 | 0.207920483 | 0.954 | 0.927 | 1.07E-14 |
| Fstl1 | 6.93E-19 | 0.089406042 | 0.03 | 0.001 | 1.30E-14 |
| Atp5g1 | 1.67E-18 | 0.219500672 | 0.916 | 0.86 | 3.13E-14 |
| Nr4a1 | 6.96E-17 | -0.292949889 | 0.124 | 0.216 | 1.30E-12 |
| Ccl3 | 7.46E-17 | -0.448042651 | 0.13 | 0.222 | 1.40E-12 |
| Rhob | 1.37E-16 | -0.347084385 | 0.384 | 0.488 | 2.56E-12 |
| Egr1 | 2.77E-16 | -0.421443561 | 0.24 | 0.321 | 5.19E-12 |
| Rplp0 | 1.65E-15 | -0.089807195 | 1 | 1 | 3.09E-11 |
| Gm19590 | 6.14E-15 | -0.314671079 | 0.226 | 0.316 | 1.15E-10 |
| Tsc22d1 | 7.07E-15 | -0.310656491 | 0.364 | 0.459 | 1.32E-10 |
| Ndufa4 | 8.04E-15 | 0.206743877 | 0.973 | 0.962 | 1.51E-10 |
| Ranbp1 | 1.17E-14 | 0.214773983 | 0.87 | 0.802 | 2.19E-10 |
| Ncl | 1.63E-14 | 0.202103566 | 0.931 | 0.896 | 3.05E-10 |
| Rpl13a | 2.96E-14 | 0.131750527 | 0.996 | 0.979 | 5.54E-10 |
| Nme1 | 3.98E-13 | 0.200117013 | 0.858 | 0.796 | 7.47E-09 |
| Prdx1 | 1.15E-12 | 0.175161926 | 0.913 | 0.868 | 2.15E-08 |
| Ypel3 | 2.11E-12 | -0.236037758 | 0.61 | 0.664 | 3.96E-08 |
| Pold4 | 2.13E-12 | -0.224562526 | 0.336 | 0.392 | 3.98E-08 |
| C1qbp | 2.16E-12 | 0.199110145 | 0.851 | 0.792 | 4.04E-08 |
| Ier3 | 2.60E-12 | -0.29883153 | 0.407 | 0.49 | 4.86E-08 |
| Jund | 3.53E-12 | -0.275725358 | 0.237 | 0.317 | 6.61E-08 |
| Hspd1 | 7.19E-12 | 0.201875098 | 0.838 | 0.789 | 1.35E-07 |
| Cdk5rap1 | 7.77E-12 | 0.149638626 | 0.107 | 0.051 | 1.46E-07 |
| Rgs2 | 9.45E-12 | -0.221462904 | 0.151 | 0.228 | 1.77E-07 |

| | | | | | |
|---|---|---|---|---|---|
| Set | 9.51E-12 | 0.204592378 | 0.788 | 0.721 | 1.78E-07 |
| Tsc22d3 | 9.99E-12 | -0.278859796 | 0.463 | 0.548 | 1.87E-07 |
| Pa2g4 | 2.12E-11 | 0.187240777 | 0.856 | 0.813 | 3.98E-07 |
| Mir142hg | 3.77E-11 | -0.249952997 | 0.52 | 0.572 | 7.06E-07 |
| Adsl | 4.01E-11 | 0.190080352 | 0.595 | 0.492 | 7.50E-07 |
| Pfn1 | 5.03E-11 | 0.098563182 | 0.995 | 0.993 | 9.43E-07 |
| Mybbp1a | 5.08E-11 | 0.191827396 | 0.611 | 0.51 | 9.51E-07 |
| Cmtm7 | 7.13E-11 | -0.138873036 | 0.98 | 0.989 | 1.34E-06 |
| Psme2 | 1.04E-10 | 0.173588482 | 0.809 | 0.743 | 1.95E-06 |
| B2m | 2.90E-10 | -0.14341061 | 0.987 | 0.988 | 5.43E-06 |
| Atp5g3 | 3.42E-10 | 0.131348508 | 0.972 | 0.962 | 6.40E-06 |
| Rps15 | 4.38E-10 | 0.140353632 | 0.929 | 0.906 | 8.20E-06 |
| Ptma | 4.54E-10 | 0.105675078 | 0.998 | 0.998 | 8.50E-06 |
| Hnrnpa2b1 | 5.10E-10 | 0.134289489 | 0.984 | 0.973 | 9.55E-06 |
| Ppid | 5.31E-10 | 0.187650452 | 0.544 | 0.447 | 9.96E-06 |
| Nhp2 | 5.74E-10 | 0.163880852 | 0.772 | 0.689 | 1.08E-05 |
| Park7 | 6.38E-10 | 0.148813447 | 0.925 | 0.907 | 1.20E-05 |
| Ldha | 7.41E-10 | 0.166904801 | 0.949 | 0.926 | 1.39E-05 |
| Cnbp | 1.03E-09 | 0.114764958 | 0.98 | 0.967 | 1.93E-05 |
| Psme1 | 1.22E-09 | 0.144818527 | 0.936 | 0.904 | 2.28E-05 |
| Rps9 | 1.30E-09 | -0.068826386 | 1 | 1 | 2.44E-05 |
| Srgn | 1.55E-09 | -0.1410578 | 0.99 | 0.992 | 2.91E-05 |
| Ppp1r15a | 1.60E-09 | -0.266025184 | 0.331 | 0.406 | 3.00E-05 |
| Camk2b | 1.62E-09 | 0.047056771 | 0.037 | 0.013 | 3.03E-05 |
| Plek | 1.65E-09 | -0.223903037 | 0.306 | 0.378 | 3.08E-05 |
| Cd69 | 1.91E-09 | -0.234200261 | 0.172 | 0.225 | 3.57E-05 |
| Snhg1 | 2.18E-09 | -0.190776198 | 0.672 | 0.708 | 4.08E-05 |
| Klhl24 | 2.71E-09 | -0.180479895 | 0.267 | 0.307 | 5.08E-05 |
| Dut | 2.73E-09 | 0.187185647 | 0.726 | 0.644 | 5.11E-05 |
| Rpl23a | 3.52E-09 | 0.170305036 | 0.862 | 0.81 | 6.59E-05 |
| Apex1 | 3.78E-09 | 0.178066408 | 0.699 | 0.615 | 7.08E-05 |
| Pnrc1 | 4.21E-09 | -0.207230777 | 0.33 | 0.401 | 7.89E-05 |
| Mat2a | 5.29E-09 | 0.158525329 | 0.712 | 0.624 | 9.91E-05 |
| Lsm7 | 5.51E-09 | 0.160651239 | 0.738 | 0.657 | 0.000103174 |
| Nolc1 | 5.61E-09 | 0.198566426 | 0.545 | 0.464 | 0.000105132 |
| Serbp1 | 5.66E-09 | 0.131466734 | 0.947 | 0.937 | 0.0001061 |
| Gart | 6.66E-09 | 0.15801399 | 0.474 | 0.382 | 0.000124691 |
| Gmfg | 6.79E-09 | -0.132373087 | 0.843 | 0.854 | 0.000127284 |
| Banf1 | 6.83E-09 | 0.150090556 | 0.889 | 0.853 | 0.000128024 |
| Lyar | 7.38E-09 | 0.158445091 | 0.455 | 0.364 | 0.000138285 |
| Nutf2 | 8.17E-09 | 0.178446845 | 0.725 | 0.65 | 0.00015312 |
| Nfkbid | 1.12E-08 | -0.183962001 | 0.133 | 0.199 | 0.000209238 |
| Txn1 | 1.19E-08 | 0.138667568 | 0.919 | 0.884 | 0.000222483 |
| Tomm6 | 1.51E-08 | 0.140531634 | 0.909 | 0.866 | 0.000282478 |

| | | | | | |
|---|---|---|---|---|---|
| Srm | 1.62E-08 | 0.201234732 | 0.462 | 0.376 | 0.000303339 |
| Timm13 | 1.65E-08 | 0.153920856 | 0.799 | 0.738 | 0.000308592 |
| Snrpd3 | 1.75E-08 | 0.143384649 | 0.811 | 0.739 | 0.000327549 |
| Tomm5 | 1.85E-08 | 0.177119054 | 0.627 | 0.551 | 0.000346637 |
| Mcm2 | 1.92E-08 | 0.153426816 | 0.509 | 0.419 | 0.000358995 |
| Itm2b | 2.00E-08 | -0.115930267 | 0.99 | 0.993 | 0.000375455 |
| Dnph1 | 2.78E-08 | 0.183791672 | 0.448 | 0.37 | 0.000520061 |
| Atp5e | 3.00E-08 | 0.119757903 | 0.954 | 0.934 | 0.000561819 |
| Tgfbr2 | 3.15E-08 | -0.169290894 | 0.179 | 0.23 | 0.000590305 |
| Eif4a1 | 3.21E-08 | 0.136219384 | 0.94 | 0.929 | 0.000600509 |
| Snrpa1 | 3.21E-08 | 0.156236379 | 0.524 | 0.435 | 0.000602098 |
| Gt(ROSA)2( | 3.50E-08 | 0.128467124 | 0.142 | 0.086 | 0.000655759 |
| Dctpp1 | 3.76E-08 | 0.177479019 | 0.758 | 0.713 | 0.000705062 |
| Vim | 4.00E-08 | -0.176991932 | 0.851 | 0.879 | 0.000750039 |
| Snrpd2 | 4.61E-08 | 0.151413674 | 0.84 | 0.793 | 0.000864613 |
| Tuba1b | 5.79E-08 | 0.184727083 | 0.553 | 0.471 | 0.001084443 |
| H2-T23 | 7.00E-08 | -0.165237876 | 0.751 | 0.789 | 0.001311541 |
| Rps17 | 7.65E-08 | 0.113583923 | 0.971 | 0.954 | 0.001432702 |
| Fus | 7.66E-08 | 0.183563641 | 0.71 | 0.648 | 0.001434529 |
| Myc | 7.74E-08 | 0.137694559 | 0.866 | 0.808 | 0.001450165 |
| Nsun2 | 9.27E-08 | 0.132610073 | 0.526 | 0.44 | 0.001737427 |
| Hspe1 | 1.08E-07 | 0.164004977 | 0.764 | 0.712 | 0.002019898 |
| Lsm2 | 1.08E-07 | 0.164725087 | 0.636 | 0.556 | 0.002022865 |
| Ruvbl2 | 1.10E-07 | 0.153558454 | 0.603 | 0.52 | 0.002066146 |
| Sf3b3 | 1.13E-07 | 0.152204842 | 0.633 | 0.55 | 0.002111321 |
| Fosb | 1.15E-07 | -0.24379067 | 0.142 | 0.199 | 0.002161048 |
| Fbl | 1.23E-07 | 0.143172925 | 0.799 | 0.731 | 0.002304656 |
| AI506816 | 1.37E-07 | 0.169135598 | 0.469 | 0.388 | 0.002572741 |
| Rps8 | 1.48E-07 | 0.048165071 | 0.999 | 1 | 0.002764796 |
| Hnrnpdl | 1.51E-07 | 0.181149887 | 0.574 | 0.499 | 0.002819886 |
| Selenop | 1.65E-07 | -0.199867746 | 0.344 | 0.395 | 0.00310042 |
| Dennd4a | 1.68E-07 | -0.193165446 | 0.344 | 0.391 | 0.003141996 |
| Syk | 2.08E-07 | -0.160790364 | 0.311 | 0.354 | 0.003891534 |
| Nop2 | 2.23E-07 | 0.161465171 | 0.373 | 0.295 | 0.00418178 |
| Eif3c | 2.29E-07 | 0.061805611 | 0.748 | 0.676 | 0.004281809 |
| Ndufc1 | 2.43E-07 | 0.14611675 | 0.56 | 0.476 | 0.004548478 |
| Plaur | 2.56E-07 | -0.19290586 | 0.182 | 0.242 | 0.004794835 |
| Rpa2 | 2.57E-07 | 0.125911932 | 0.494 | 0.413 | 0.00481925 |
| Gas5 | 2.58E-07 | -0.105935818 | 0.987 | 0.985 | 0.004825084 |
| H3f3b | 2.65E-07 | -0.114093137 | 0.995 | 0.996 | 0.004966849 |
| Mrpl12 | 2.70E-07 | 0.159295917 | 0.715 | 0.655 | 0.0050501 |
| Mrpl17 | 2.78E-07 | 0.127575 | 0.631 | 0.549 | 0.005213334 |
| Tpi1 | 2.78E-07 | 0.134881696 | 0.83 | 0.772 | 0.005214848 |
| U2af1 | 2.90E-07 | 0.158679942 | 0.774 | 0.725 | 0.005433654 |

| | | | | | |
|---|---|---|---|---|---|
| Nop56 | 2.91E-07 | 0.172271907 | 0.579 | 0.504 | 0.005457363 |
| Paip2 | 3.24E-07 | -0.131493487 | 0.77 | 0.778 | 0.006067175 |
| Ybx1 | 3.76E-07 | 0.156933836 | 0.687 | 0.616 | 0.007047033 |
| Pdia6 | 4.05E-07 | 0.108560297 | 0.705 | 0.627 | 0.007580451 |
| Tbc1d9b | 4.08E-07 | -0.166445243 | 0.278 | 0.327 | 0.007645681 |
| Ddx39 | 4.30E-07 | 0.152646497 | 0.517 | 0.435 | 0.00806378 |
| Rexo2 | 4.35E-07 | 0.140844267 | 0.599 | 0.518 | 0.008145568 |
| Mef2c | 4.62E-07 | -0.190791735 | 0.539 | 0.592 | 0.008650697 |
| Rabgap1l | 4.97E-07 | -0.10761435 | 0.465 | 0.462 | 0.009303767 |
| Ssb | 5.40E-07 | 0.147640907 | 0.765 | 0.708 | 0.010120599 |
| Mrto4 | 5.48E-07 | 0.07376865 | 0.502 | 0.432 | 0.01026955 |
| Alad | 6.12E-07 | 0.109170782 | 0.423 | 0.347 | 0.011461349 |
| Akap13 | 6.63E-07 | -0.182661038 | 0.729 | 0.766 | 0.012420623 |
| mt-Co3 | 7.32E-07 | -0.132112581 | 0.98 | 0.983 | 0.013718232 |
| Eif3b | 7.42E-07 | 0.154967083 | 0.608 | 0.531 | 0.013902995 |
| Vps28 | 7.44E-07 | 0.094462701 | 0.65 | 0.574 | 0.013942368 |
| Hint1 | 7.58E-07 | 0.109826003 | 0.967 | 0.959 | 0.014201408 |
| Ddit4 | 7.97E-07 | -0.199582876 | 0.229 | 0.292 | 0.014939624 |
| Nufip1 | 8.21E-07 | 0.10652751 | 0.295 | 0.229 | 0.015372541 |
| Ipo5 | 8.39E-07 | 0.157615003 | 0.416 | 0.34 | 0.015717025 |
| Nme2 | 8.54E-07 | 0.094628619 | 0.979 | 0.974 | 0.016000652 |
| Dnajc9 | 8.73E-07 | 0.146047234 | 0.458 | 0.38 | 0.016359291 |
| Lyz2 | 9.08E-07 | -0.130456475 | 0.66 | 0.672 | 0.017013991 |
| Got2 | 9.59E-07 | 0.117033081 | 0.452 | 0.375 | 0.017968214 |
| Dancr | 1.02E-06 | 0.168459701 | 0.302 | 0.234 | 0.019117823 |
| Pdcd4 | 1.03E-06 | -0.154149982 | 0.329 | 0.367 | 0.019222173 |
| Dapp1 | 1.16E-06 | -0.160286323 | 0.656 | 0.683 | 0.021669555 |
| Gimap6 | 1.16E-06 | -0.204288857 | 0.387 | 0.437 | 0.021788675 |
| St6galnac4 | 1.23E-06 | 0.101313462 | 0.321 | 0.253 | 0.023135204 |
| Cox5a | 1.26E-06 | 0.12093324 | 0.922 | 0.895 | 0.023626026 |
| Prmt1 | 1.27E-06 | 0.1360837 | 0.81 | 0.749 | 0.023804857 |
| Atic | 1.39E-06 | 0.152250592 | 0.606 | 0.535 | 0.026094953 |
| Dnajb1 | 1.44E-06 | -0.138695811 | 0.458 | 0.483 | 0.027062982 |
| Nop58 | 1.72E-06 | 0.056536721 | 0.764 | 0.698 | 0.032248016 |
| Timm17a | 1.74E-06 | 0.023483234 | 0.415 | 0.366 | 0.032641871 |
| Mgat2 | 2.02E-06 | 0.140336594 | 0.471 | 0.395 | 0.037810452 |
| Cox6c | 2.04E-06 | 0.113899434 | 0.9 | 0.856 | 0.038293515 |
| Rnaseh2c | 2.16E-06 | 0.143793075 | 0.61 | 0.535 | 0.040483307 |
| AC149090. | 2.17E-06 | -0.162165418 | 0.497 | 0.539 | 0.040725289 |
| Hsp90aa1 | 2.23E-06 | 0.119952039 | 0.947 | 0.934 | 0.041703601 |
| Per1 | 2.32E-06 | -0.156549261 | 0.224 | 0.276 | 0.043436058 |
| Nudc | 2.40E-06 | 0.14516303 | 0.797 | 0.748 | 0.044901413 |
| Rpl5 | 2.43E-06 | 0.097152481 | 0.96 | 0.951 | 0.045460681 |
| Ripor1 | 2.45E-06 | -0.159522808 | 0.5 | 0.529 | 0.045928258 |

| | | | | | |
|---|---|---|---|---|---|
| Gnl3 | 2.46E-06 | 0.103969529 | 0.426 | 0.354 | 0.046001258 |
| Tuba4a | 2.54E-06 | 0.108508862 | 0.565 | 0.489 | 0.047624931 |
| Fkbp11 | 2.61E-06 | 0.144614258 | 0.203 | 0.145 | 0.048976845 |
| Comtd1 | 2.65E-06 | 0.125360191 | 0.361 | 0.289 | 0.049580126 |

**Supplemental Table III - DEGs from Proliferating cell cluster**

| | p_val | avg_log2FC | pct.1 | pct.2 | p_val_adj |
|---|---|---|---|---|---|
| Jun | 3.83E-37 | -0.703596694 | 0.702 | 0.838 | 7.18E-33 |
| Mif | 1.13E-22 | 0.302480297 | 0.991 | 0.982 | 2.12E-18 |
| Zfp36 | 3.28E-22 | -0.494000526 | 0.684 | 0.799 | 6.15E-18 |
| Fstl1 | 1.75E-18 | 0.080551696 | 0.052 | 0.001 | 3.28E-14 |
| Fos | 1.31E-15 | -0.499733051 | 0.669 | 0.75 | 2.46E-11 |
| Ypel3 | 4.71E-15 | -0.345367006 | 0.511 | 0.603 | 8.83E-11 |
| Dusp1 | 1.68E-14 | -0.405673575 | 0.422 | 0.567 | 3.16E-10 |
| Junb | 1.61E-13 | -0.326973024 | 0.637 | 0.725 | 3.01E-09 |
| Ppia | 1.73E-13 | 0.105247524 | 1 | 1 | 3.25E-09 |
| Ier2 | 2.21E-13 | -0.348792242 | 0.697 | 0.794 | 4.14E-09 |
| Ranbp1 | 3.50E-13 | 0.246474455 | 0.963 | 0.938 | 6.55E-09 |
| Ldha | 3.75E-13 | 0.226489693 | 0.985 | 0.971 | 7.02E-09 |
| Rps15 | 5.64E-13 | 0.174713425 | 0.989 | 0.979 | 1.06E-08 |
| 2410006H16R | 1.63E-12 | -0.27900707 | 0.923 | 0.942 | 3.06E-08 |
| Nme1 | 2.48E-12 | 0.213867292 | 0.955 | 0.915 | 4.64E-08 |
| Egr1 | 4.42E-12 | -0.377293775 | 0.254 | 0.372 | 8.27E-08 |
| Pa2g4 | 5.04E-12 | 0.208301865 | 0.973 | 0.94 | 9.43E-08 |
| Dctpp1 | 6.89E-12 | 0.233631735 | 0.933 | 0.855 | 1.29E-07 |
| Atp5g3 | 1.58E-11 | 0.143411568 | 0.999 | 0.995 | 2.96E-07 |
| Atp5g1 | 1.85E-11 | 0.179890193 | 0.982 | 0.967 | 3.47E-07 |
| Hspd1 | 3.09E-11 | 0.223382618 | 0.935 | 0.879 | 5.79E-07 |
| Shmt2 | 3.59E-11 | 0.204866059 | 0.715 | 0.582 | 6.73E-07 |
| mt-Co1 | 3.66E-11 | -0.176898453 | 1 | 1 | 6.85E-07 |
| Dut | 3.73E-11 | 0.225520054 | 0.944 | 0.872 | 6.99E-07 |
| Tpi1 | 3.89E-11 | 0.200544012 | 0.953 | 0.908 | 7.28E-07 |
| Prdx1 | 4.14E-11 | 0.162493429 | 0.991 | 0.975 | 7.75E-07 |
| C1qbp | 6.29E-11 | 0.201119373 | 0.952 | 0.912 | 1.18E-06 |
| Hsp90aa1 | 6.90E-11 | 0.154825321 | 0.998 | 0.991 | 1.29E-06 |
| Rgs1 | 1.51E-10 | -0.33487413 | 0.356 | 0.482 | 2.83E-06 |
| Rpl5 | 1.72E-10 | 0.147715316 | 0.995 | 0.984 | 3.22E-06 |
| Fbl | 2.60E-10 | 0.186350866 | 0.931 | 0.879 | 4.87E-06 |
| Ran | 2.64E-10 | 0.135534588 | 1 | 0.999 | 4.95E-06 |
| Btg2 | 3.47E-10 | -0.313753926 | 0.391 | 0.485 | 6.51E-06 |
| Nr4a1 | 4.56E-10 | -0.211196591 | 0.154 | 0.252 | 8.55E-06 |
| Ccl4 | 4.84E-10 | -0.305640736 | 0.137 | 0.229 | 9.07E-06 |
| Rhob | 1.20E-09 | -0.239811387 | 0.316 | 0.439 | 2.25E-05 |
| Set | 1.95E-09 | 0.220090521 | 0.902 | 0.845 | 3.65E-05 |

| | | | | | |
|---|---|---|---|---|---|
| Tomm40 | 2.74E-09 | 0.201783779 | 0.804 | 0.702 | 5.14E-05 |
| Ctla2a | 3.87E-09 | -0.304394287 | 0.89 | 0.919 | 7.25E-05 |
| Eif4a1 | 5.04E-09 | 0.151918407 | 0.991 | 0.984 | 9.44E-05 |
| Ncl | 5.09E-09 | 0.201387801 | 0.986 | 0.963 | 9.53E-05 |
| Pde4b | 5.29E-09 | -0.263302578 | 0.717 | 0.78 | 9.92E-05 |
| Nop10 | 7.69E-09 | 0.148893564 | 0.986 | 0.967 | 0.000144057 |
| Cdca7 | 1.07E-08 | 0.171357344 | 0.752 | 0.634 | 0.000200964 |
| Ptma | 1.10E-08 | 0.085038975 | 1 | 1 | 0.000205572 |
| Cox5a | 1.18E-08 | 0.127933961 | 0.988 | 0.988 | 0.000220687 |
| Timm8a1 | 1.23E-08 | 0.19582392 | 0.703 | 0.591 | 0.000230139 |
| Mrto4 | 1.51E-08 | 0.177151637 | 0.74 | 0.625 | 0.000282386 |
| Cnbp | 2.01E-08 | 0.130110174 | 0.994 | 0.991 | 0.000375742 |
| H2afv | 2.02E-08 | -0.178599457 | 0.995 | 0.993 | 0.000377583 |
| Slc25a5 | 2.54E-08 | 0.145955903 | 0.988 | 0.978 | 0.000475941 |
| Selenow | 2.74E-08 | 0.159275663 | 0.974 | 0.944 | 0.000512506 |
| Klf6 | 2.78E-08 | -0.244085312 | 0.257 | 0.356 | 0.000521736 |
| Eif5a | 3.37E-08 | 0.12165318 | 0.999 | 0.998 | 0.00063203 |
| Hspe1 | 4.54E-08 | 0.171034229 | 0.916 | 0.852 | 0.000850853 |
| Tmem223 | 4.60E-08 | 0.188048913 | 0.761 | 0.658 | 0.000861263 |
| Cacybp | 9.33E-08 | 0.144623699 | 0.909 | 0.834 | 0.001748188 |
| Nutf2 | 1.07E-07 | 0.156490644 | 0.916 | 0.86 | 0.002005255 |
| Ier3 | 1.19E-07 | -0.253427239 | 0.439 | 0.491 | 0.00222644 |
| Cdc20 | 1.23E-07 | -0.269666201 | 0.701 | 0.742 | 0.002303849 |
| Cenpa | 1.32E-07 | -0.276695911 | 0.688 | 0.779 | 0.002476228 |
| Polr2f | 1.47E-07 | 0.180138841 | 0.859 | 0.798 | 0.002744975 |
| Mat2a | 1.60E-07 | 0.189008321 | 0.811 | 0.726 | 0.003004895 |
| Park7 | 1.88E-07 | 0.128257269 | 0.989 | 0.983 | 0.003531291 |
| Pfn1 | 1.95E-07 | 0.101581051 | 0.999 | 0.999 | 0.00364988 |
| Itm2b | 2.00E-07 | -0.133443914 | 0.996 | 0.998 | 0.003749475 |
| Aprt | 2.03E-07 | 0.174805966 | 0.871 | 0.798 | 0.00379803 |
| Zfp36l2 | 2.46E-07 | -0.211756269 | 0.792 | 0.824 | 0.004605973 |
| Rpl31 | 2.63E-07 | 0.118373658 | 0.996 | 0.984 | 0.004924692 |
| Srm | 2.69E-07 | 0.170246928 | 0.574 | 0.459 | 0.005044747 |
| Atp5e | 3.05E-07 | 0.123511752 | 0.991 | 0.987 | 0.005716948 |
| Nme2 | 3.31E-07 | 0.126636911 | 0.998 | 0.991 | 0.006195478 |
| Apex1 | 3.94E-07 | 0.124973785 | 0.819 | 0.722 | 0.007389524 |
| Ndufb6 | 4.11E-07 | 0.147270732 | 0.89 | 0.825 | 0.00769366 |
| Tsc22d3 | 6.24E-07 | -0.219503647 | 0.509 | 0.59 | 0.011691833 |
| Mrpl12 | 6.33E-07 | 0.136482674 | 0.909 | 0.839 | 0.011866714 |

| | | | | | |
|---|---|---|---|---|---|
| Ccl3 | 7.02E-07 | -0.162040546 | 0.072 | 0.128 | 0.013143575 |
| Pimreg | 7.61E-07 | -0.218622696 | 0.461 | 0.535 | 0.014260195 |
| Cks2 | 8.25E-07 | -0.205389549 | 0.966 | 0.966 | 0.015448883 |
| Gmfg | 1.05E-06 | -0.172948117 | 0.918 | 0.939 | 0.019677876 |
| AI506816 | 1.12E-06 | 0.16949006 | 0.784 | 0.697 | 0.020921326 |
| Arl6ip1 | 1.21E-06 | -0.237522039 | 0.944 | 0.941 | 0.022754515 |
| Sumo2 | 1.63E-06 | 0.113504222 | 0.993 | 0.992 | 0.030532357 |
| Rps6 | 1.80E-06 | 0.078097486 | 1 | 1 | 0.033764772 |
| Rpl18a | 1.96E-06 | -0.063560182 | 1 | 1 | 0.036730229 |
| Nsa2 | 2.08E-06 | -0.151558585 | 0.891 | 0.918 | 0.038887313 |
| Rplp0 | 2.37E-06 | -0.067580809 | 1 | 1 | 0.044411948 |
| Gm42047 | 2.51E-06 | -0.171147316 | 0.195 | 0.256 | 0.047069836 |
| Vim | 2.62E-06 | -0.21020195 | 0.853 | 0.875 | 0.04913578 |
